# Systematic comparison of high-throughput single-cell RNA-seq methods for immune cell profiling

**DOI:** 10.1101/2020.09.04.283499

**Authors:** Tracy M. Yamawaki, Daniel R. Lu, Daniel C. Ellwanger, Dev Bhatt, Paolo Manzanillo, Hong Zhou, Oh Kyu Yoon, Oliver Homann, Songli Wang, Chi-Ming Li

**Affiliations:** Genome Analysis Unit, Amgen Research, South San Francisco, California, United States of America; Oncology/Inflammation, Amgen Research, South San Francisco, California, United States of America

**Keywords:** Single cell, Transcriptomics, single-cell RNA-seq, High Throughput Sequencing, immune-cell profiling

## Abstract

**Background:** Elucidation of immune populations with single-cell RNA-seq has greatly benefited the field of immunology by deepening the characterization of immune heterogeneity and leading to the discovery of new subtypes. However, single-cell methods inherently suffer from limitations in the recovery of complete transcriptomes due to the prevalence of cellular and transcriptional dropout events. This issue is often compounded by limited sample availability and limited prior knowledge of heterogeneity, which can confound data interpretation.

**Results:** Here, we systematically benchmarked seven high-throughput single-cell RNA-seq methods. We prepared 21 libraries under identical conditions of a defined mixture of two human and two murine lymphocyte cell lines, simulating heterogeneity across immune-cell types and cell sizes. We evaluate methods by their cell recovery rate, library efficiency, sensitivity, and ability to recover expression signatures for each cell type. We observed higher mRNA detection sensitivity with the 10x Genomics 5’ v1 and 3’ v3 methods. We demonstrate that these methods have fewer drop-out events which facilitates the identification of differentially-expressed genes and improves the concordance of single-cell profiles to immune bulk RNA-seq signatures.

**Conclusion:** Overall, our characterization of immune cell mixtures provides useful metrics, which can guide selection of a high-throughput single-cell RNA-seq method for profiling more complex immune-cell heterogeneity usually found *in vivo*.

## Background

Understanding the cellular diversity underlying immune responses is an important component of immunological research. Techniques such as FACS and mass cytometry [1] are useful for studying cellular diversity according to well-characterized cell-surface-protein markers. The advent of single-cell RNA-seq has expanded the power to characterize individual immune cells from a defined set of cell-surface markers to the entire transcriptome. These single-cell technologies have enabled immunologists to characterize inflammation [1] and immune responses to cancer [1-2], uncovering previously uncharacterized cellular diversity and cell-type specific transcriptional responses. As recent advances have increased cell throughput and lowered per-cell costs, the number of high-throughput techniques that can process more than a thousand cells per experiment has increased.

Several key factors, such as variable capture and amplification efficiencies during library preparation, impact the ability of single-cell RNA-seq techniques to accurately and comprehensively characterize immune-cell diversity. Mixtures of different cell sizes are particularly complex as small cells contain low total number of transcripts and therefore, are difficult to distinguish from ambient noise.

The relatively small size and low mRNA content of immune cells may impact the performance of single-cell RNA-seq methods differently than was previously described using larger cells [1-2]. Immune cells constitute a broad range of cell types across various lineages, activation states, and cell sizes. Efficient recovery across these diverse cell types impacts the fidelity of cell-composition analyses. Methods that recover a larger fraction of cells in a cost-efficient manner benefit studies that sample tissues containing few immune cells. Also, increased sensitivity in detecting individual mRNA transcripts results in more comprehensive cellular profiles, which greatly advances the characterization of immune sub-types. A more complete picture of cellular transcriptional activity facilitates the identification of differentially expressed (DE) marker genes and positively impacts the mapping of cells against reference immune cell signatures.

Previous benchmarking studies using somatic cell lines or peripheral blood mononuclear cells (PBMCs) reported that high-throughput single-cell RNA-seq methods generally enable broader sampling of diverse populations at a lower per-cell cost. However, larger sample sizes come at the expense of lower mRNA detection sensitivity [1-2]. In this work, we extend previous findings with a focus on the application of high-throughput methods to immune-cell profiling. By using a defined mixture of four lymphocyte cell lines, we assess the performance of seven high-throughput methods using four commercially-available systems to address common concerns in immune-cell profiling. First, we examine library efficiency in terms of cell recovery and cell-assignable reads. Next, we assess mRNA detection sensitivity and the correlation of cellular profiles to immune cell signatures from bulk RNA-seq. Finally, we compare results across cell types in consideration of varying cell sizes and cellular mRNA contents. This study serves as useful guidelines for the selection of a suitable single-cell RNA-seq method to study immune cells.

## Results

### Design of Single-cell RNA-seq Benchmarking Experiments

We benchmarked four commercially-available high-throughput single-cell systems: the Chromium (10x Genomics) [1], the ddSEQ (Illumina and Bio-Rad), the scRNA-Seq System running Drop-seq (Dolomite Bio) [1], and the ICELL8 cx (Takara Bio) [1] (**Figure 1**). We tested three methods available for the Chromium (3’ v3, 3’ v2 and 5’ v1) as well as two methods for the ICELL8 (the official 3’ DE protocol and an alternate 3’ DE-UMI protocol). All methods tested perform mRNA end counting by tagging mRNA sequences with a barcode containing a cell identifier (CID) and a unique molecular identifier (UMI) with lengths that vary by method (**Supplemental Table 1)**.

**Figure 1:**
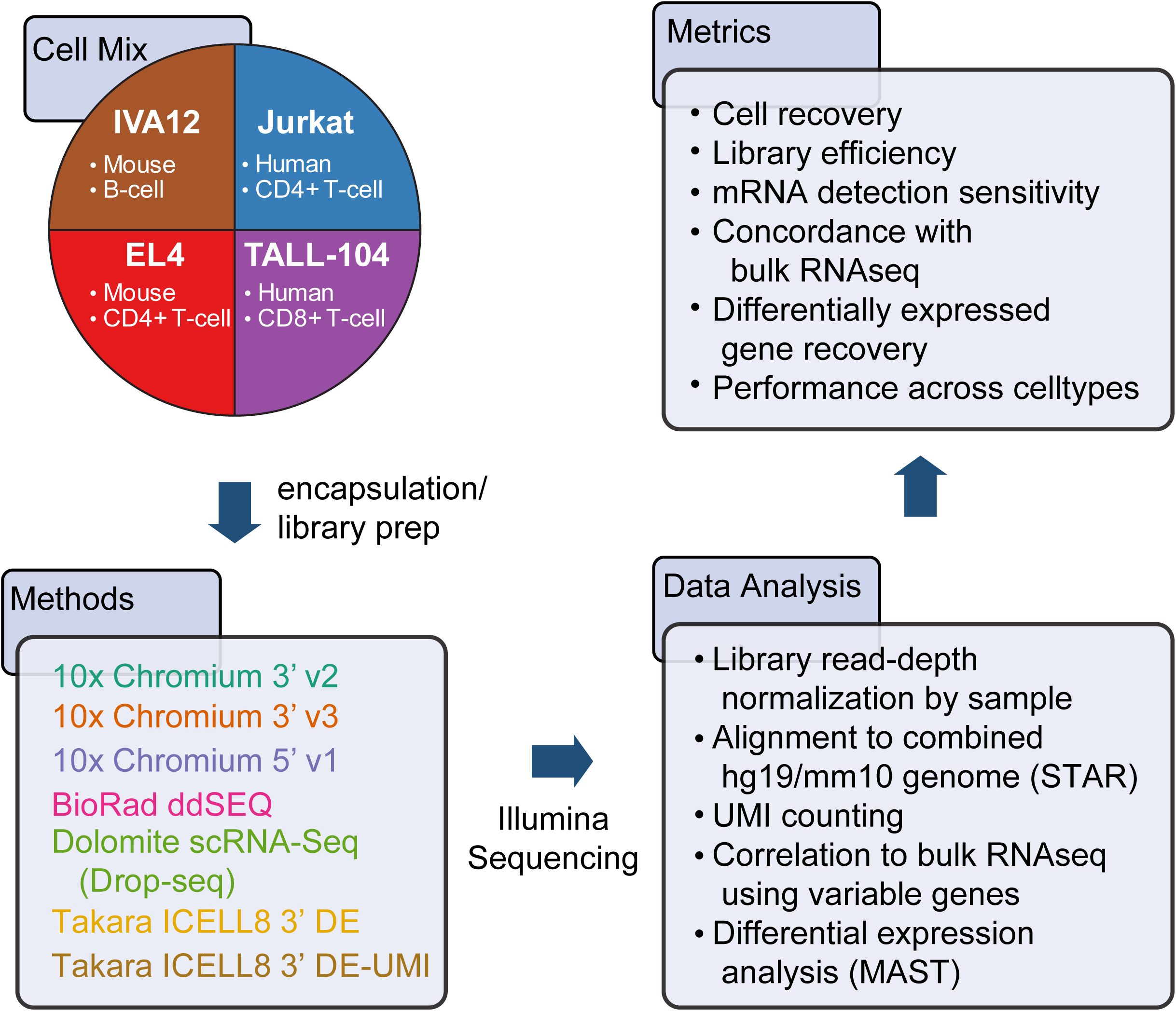
Overview of high-throughput single-cell benchmarking experiments. Experiments were performed using four immune cell lines to benchmark cell recovery, transcript detection sensitivity, concordance to bulk RNA-seq and differentially-expressed gene identification.

All techniques, apart from ddSEQ, amplify full-length cDNA (**Supplemental Table 1**) using a modified Smart-seq protocol [1, 2], which incorporates a 5’ PCR handle by employing a reverse transcriptase’s ability to switch templates at the end of a transcript. Full-length cDNA can be amplified with primers in the 5’ template-switch and 3’ poly-T oligonucleotides. Barcoded cDNA ends are further amplified after direct ligation or tagmentation to incorporate Illumina sequencing adapters. ddSEQ contains a single amplification step during adapter incorporation after second strand synthesis without amplification of full-length cDNA. Amplification bias introduced in the multiple rounds of PCR in these protocols, is mitigated by the incorporation of UMIs [1]. However, UMI counts are unreliable in the ICELL8 3’ DE protocol because cDNA is amplified in the presence of barcoding primers, potentially inflating UMI counts. The alternative ICELL8 3’ DE-UMI protocol is more robust for UMI counting since reverse transcription and cDNA amplification are uncoupled by an exonuclease digestion of barcoding primers.

We used a 1:1:1:1 mixture of four lymphocyte cell lines from two species (**Figure 1 and Supplemental Table 2**): EL4 (mouse CD4+ T cells), IVA12 (mouse B cells), Jurkat (human CD4+ T cells), and TALL-104 (human CD8+ T cells). These cells also vary in morphology: TALL-104 cells (∼5 µm diameter) are considerably smaller than the other cell types (∼10 µm diameter). These cell lines are expected to have distinct expression profiles enabling the classification of each cell type. Usage of cells from two species allowed us to clearly identify cross-species doublet contamination to calculate capture rates of cell multiplets. To mirror typical single-cell sequencing runs and to ensure a comparison independent of sequencing limitations, we normalized the read depth of our libraries to ∼50,000 reads per cell (**Figure 1, Supplemental Figure 1 and 2**). Cells were identified and classified by correlating single-cell expression profiles to bulk RNA-seq.

**Figure 2:**
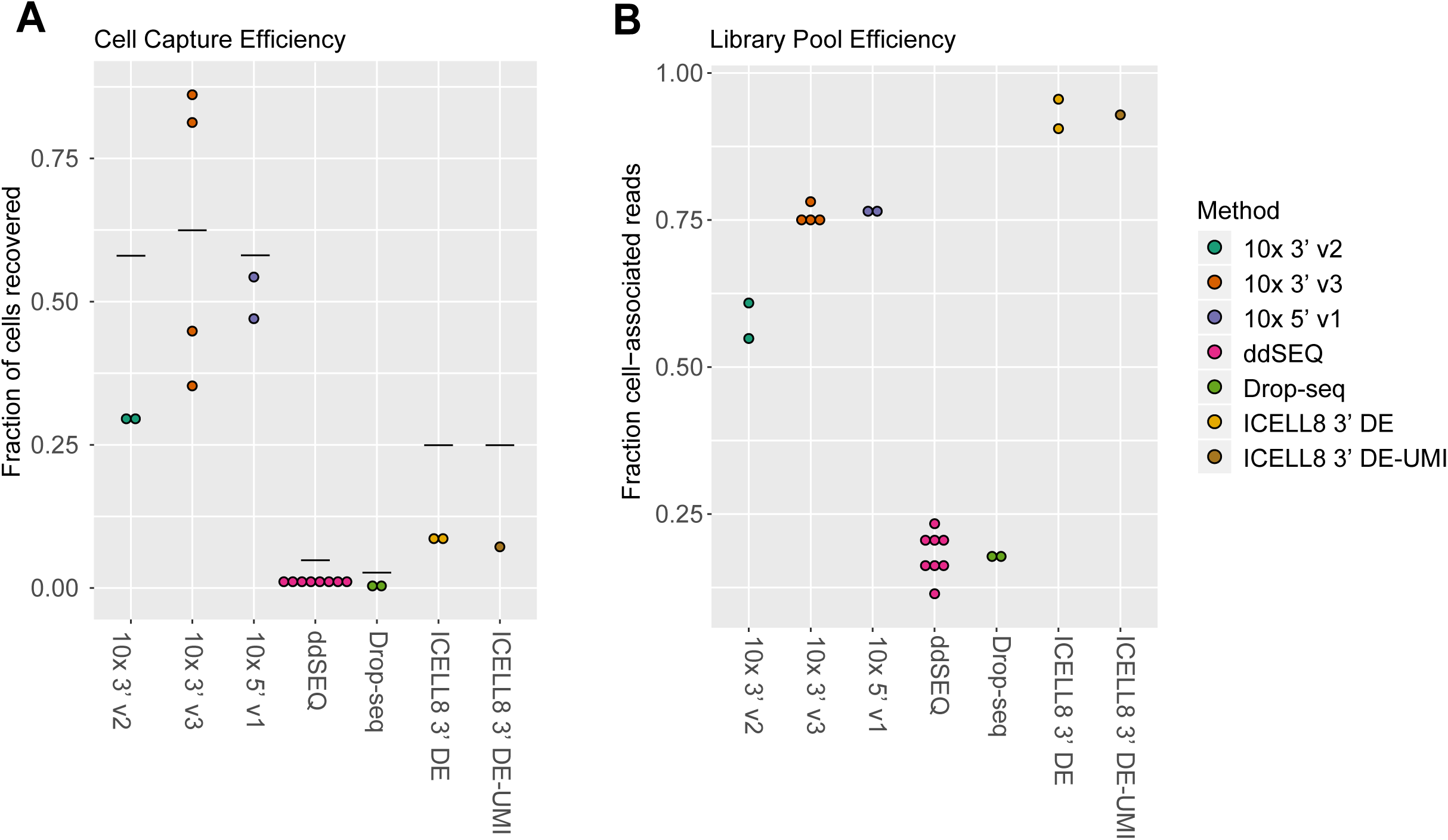
Library-pool and cell-capture efficiencies. (A) Cell capture efficiency was measured by the number of cell identifiers (CIDs) above the inflection point of the rank ordered reads/CID plot (knee plot) relative to the number of cells loaded on the instrument. Horizontal lines indicate theoretical capture efficiency based on bead/cell loading concentrations or manufacturer’s guidelines. (B) Library pool efficiency was measured by the number of reads in CIDs above the inflection point.

### Evaluation of Cell Capture and Library Efficiency

One important consideration for single-cell RNA-seq is the capture rate, or the fraction of cells recovered in the data relative to input. This is especially critical when working with precious samples with few cells. To identify recovered cells, we used the curve of the log-total count against the log-rank of each CID, which is equivalent to the transposed log-log empirical cumulative density plot of the total counts of each CID. The knee and inflection points in this curve typically define the transition between the cell-containing component and the ambient RNA component of the total count distribution. Here, we defined a recovered cell as a CID located above the inflection point (**Supplemental Figure 2A)**. In our tests, we find that capture rates are slightly lower than, but track with theoretical rates (**Figure 2A and Table 1**). As expected, we observed the highest rates with 10x Genomics methods, ranging from ∼30 to ∼80%, while ddSEQ and Drop-seq methods recovered < 2% of cells.

**Table 1:** Summary of average mRNA/gene detection sensitivities and capture rates for each method.

| Method | Avg Multiplet Rate | Avg Cell Capture Efficiency | Avg Library Pool Efficiency | Median nUMIs (EL4) | Median nGenes (EL4) | GD<sub>50</sub> (EL4) | Avg nDE genes | Avg nDE genes (> 1.5 FC in bulk) |
| --- | --- | --- | --- | --- | --- | --- | --- | --- |
| 10x 3' v2 | 0.46% | 29.50% | 57.90% | 21,570 | 3,882 | 20.2 FPKM | 3,314 | 2,711 |
| 10x 3' v3 | 1.75% | 61.90% | 75.90% | 28,006 | 4,776 | 13.6 FPKM | 4,005 | 3,388 |
| 10x 5' v1 | 0.49% | 50.70% | 76.50% | 25,988 | 4,470 | 16.8 FPKM | 4,797 | 3,491 |
| ddSEQ | 0.45% | 1.01% | 18.10% | 10,466 | 3,644 | 25 FPKM | 2,740 | 2,397 |
| Drop-seq | 0.55% | 0.36% | 17.80% | 8,791 | 3,255 | 26.7 FPKM | 2,824 | 2,504 |
| ICELL8 3' DE | 2.18% | 8.63% | 93.00% | 16,909 | 2,849 | 37.9 FPKM | 1,815 | 1,528 |
| ICELL8 3' DE-UMI | 0.98% | 7.20% | 92.90% | 2,792 | 1,288 | 112.1 FPKM | 985 | 861 |

\*: The value with the best performance for each parameter is highlighted

In addition to the capture rate, we also quantified events capturing multiple cells in a single partition. This technical artifact impairs downstream data analysis, as artificial mixtures of transcriptomes may be interpreted wrongly as single cells. The extent of this issue is influenced by the quality of the single-cell suspension, cell health, and cell loading concentration. By counting CIDs with a significant fraction of both human and mouse transcripts, for all methods, we observed multiplet rates around the 5% we had targeted with our cell-loading concentrations (**Supplemental Figure 3A and Table 1**).

Another significant factor in efficiency is the fraction of reads that can be assigned to individual cells. Increased background noise in sequencing libraries results in wasted reads and unnecessarily increased sequencing costs. We observed the highest fraction of cell-associated reads for our ICELL8 experiments (> 90%), intermediate rates for 10x experiments (∼50-75%) and the lowest rates for ddSEQ and Drop-seq (< 25%) (**Figure 2B and Supplemental Tables 3 & 4**). We also examined the genomic locations of aligned reads. About 75% of aligned bases of each library were mapped to exons and UTRs. Notably, the intergenic fraction was lowest in 10x samples, suggesting lower genomic contamination in these methods. (**Supplemental Figure 3B**). The ddSEQ method exhibited the greatest UTR bias. This is likely due to the longest read-length (150 bases) for ddSEQ of each tested technology.

### 10x 5’ v1 and 3’ v3 Methods Demonstrate the Highest mRNA Detection Sensitivity

Because immune cells tend to have low levels of mRNA, the mRNA detection sensitivity, or the fraction of a cell’s transcriptome detectable, critically impacts downstream analyses. Single-cell RNA-seq methods are inherently prone to dropouts due to inefficiencies during library preparation resulting in false negative gene expression signals [1]. Although we performed library normalization to obtain a consistent read depth across all cells, we found that read distributions of individual cell types varied.

Since EL4 cells demonstrated the highest consistency between read distributions across experiments (**Supplemental Figure 1C**), we focused our initial analysis on EL4 cells to minimize batch effects due to differential sequencing depths. We observed the highest detection of both transcripts and genes with at least one read count using 10x Genomics methods, with the highest levels seen in the 3’ v3 experiments (median 28,006 UMIs/4,776 genes across all samples) followed by the 5’ v1 and 3’ v2 kits (25,988 UMIs/4,470 genes and 21,570 UMIs/3,882 genes, respectively) (**Figure 3A-B and Supplemental Table 4**). ddSEQ and Drop-seq experiments demonstrated similar detection rates (10,466 UMIs/3,644 genes and 8,791 UMIs/3,255 genes respectively). UMI counts generated by the ICELL8 3’ DE method are unreliable due to residual barcoding primers during cDNA amplification, so we focused on gene detection sensitivity instead. We observed a significant drop in gene detection between the 3’ DE and 3’ DE-UMI methods (2,849 and 1,288 genes respectively) and a low number of UMIs counted in the 3’ DE-UMI method (2,792 UMIs). This suggests that many transcripts are lost in the additional primer digestion and cleanup steps. For the other three cell types, rankings of methods by absolute UMI-and gene-count distributions slightly differed from EL4 cells, likely due to greater variation in read depth across samples for these cell types (**Supplemental Figures 1C and 4A**).

**Figure 3:**
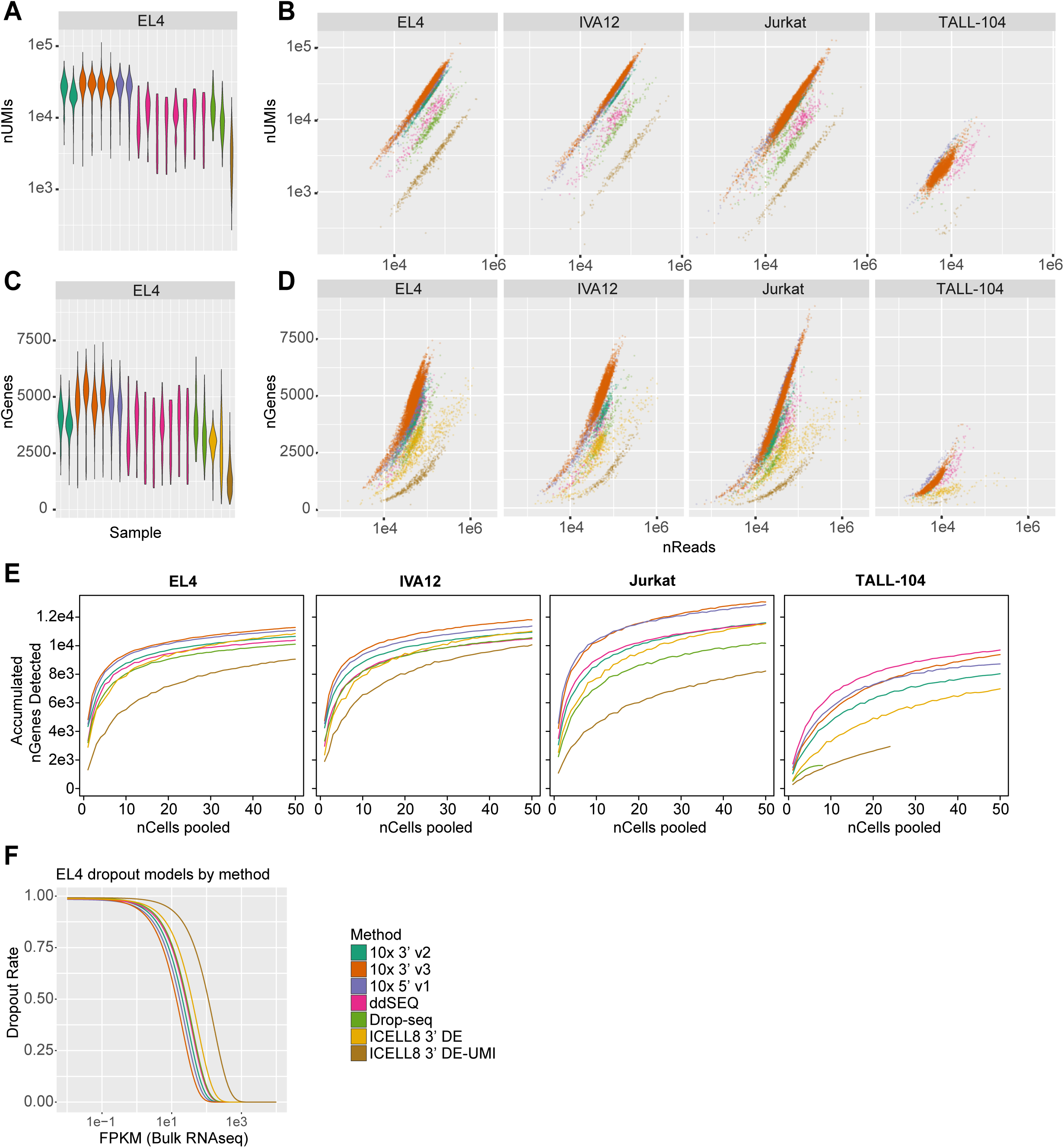
Transcript detection sensitivity. (A) Distributions of unique molecular identifiers (UMIs) detected in EL4 cells by sample are plotted. (B) Numbers of UMIs detected versus numbers of reads per cell for each cell type are plotted. (C) Distributions of genes detected in EL4 cells by sample are plotted. (D) Numbers of genes detected versus numbers of reads per cell are plotted. (E) Average number of genes detected from aggregated data of subsamples up to 50 cells was plotted. (F) Dropout modeling (dropout rate versus FPKM of bulk sequencing) for EL4 cells by method are shown. A left-shifted curve indicates higher sensitivity, that is, fewer dropouts at lower expression levels. Sensitivity of methods for EL4 cells ranked in the following order: 10x 3’ v3 > 10x 5’ v1 > 10x 3’ v2 > ddSEQ > Drop-seq > ICELL8 3’ DE > ICELL8 3’ DE-UMI. Cells with high mitochondrial expression rates were excluded from this calculation.

To account for varying read distributions across the four cell types (**Supplemental Figure 1C**), we compared the number of detected UMIs and genes relative to the total number of reads per cell. For EL4, IVA12 and Jurkat cells, we see a similar trend across methods with regards to efficiency of transcript and gene detection (**Figure 3B & 3D**). Again, 10x 3’ v3 (mean±SD reads/UMI = 2.07 ± 0.52, reads/gene = 9.04 ± 2.65) and 5’ v1 chemistries (mean±SD reads/UMI = 1.98 ± 0.19, reads/gene = 9.51 ± 2.68) were the most efficient, requiring fewer reads to detect a single UMI or gene. These methods are followed by 10x 3’ v2 (reads/UMI = 2.35 ± 0.33, reads/gene = 11.17 ± 3.03), ddSEQ (reads/UMI = 5.25 ± 1.14, reads/gene = 13.42 ± 3.89), Drop-seq (reads/UMI = 6.40 ± 1.42, reads/gene = 15.97 ± 5.62) and ICELL8 methods (3’ DE: reads/gene = 29.68 ± 41.48, 3’DE-UMI: reads/UMI = 21.77 ± 5.50, reads/gene = 47.5 ± 17.91). This trend is largely mirrored in TALL-104 cells, albeit less distinct due to the low read depth obtained for those cells (**Figure 3B-C and Supplemental Figure 1C**).

We further examined the number of genes with at least one sequenced read in pseudo-bulk populations. For this purpose, cells form each cell type were pooled and gene expression measurements were merged. We observed similar trends with higher numbers of detected genes with the 10x 3’ v3, and 5’ v1 method for EL4, IVA12 and Jurkat cells (**Figure 3E**). Although the ICELL8 3’ DE method had a low per-cell gene detection rate, when pooling more than 30 cells this method exhibited comparable levels of gene detection to 10x 3’ v2, ddSEQ and Drop-seq methods. This is likely due to the high false negative rate of genes with overall low expression levels in the ICELL8 3’ DE method. The cumulative number of genes for TALL-104 cells was lower than the other cell types and the relative detection rates across methods did match trends seen in other cell types, possibly due to the low read depth and cell recovery for this cell type.

We also examined the ability of each method to detect genes at various expression levels by calculating the dropout rate, the conditional probability that a gene is not detected in a given cell. The dropout rate was modeled as a function of the expression level in bulk RNA-seq (FPKM) for each cell type. We used a nonlinear least square fit of the data that accounts for the activity of reverse transcriptase described by Michaelis-Menten kinetics [1-2]. Here, higher gene detection sensitivity as a function of fewer dropouts at lower expression levels, is indicated by left-shifted curves and lower Gene Detection 50 (GD_50_) value, the point at which this curve reaches a detection probability of 0.5. The GD_50_ metric represents the expression level of a gene we would expect to be detected in half of the cells, and can help guide expectations of detection rates for genes of interest based on their expression in bulk RNA-seq. For EL4 cells, 10x Genomics methods were the most sensitive with 10x 3’ v3 having the lowest GD_50_ at 13.6 FPKM, followed by the 5’ v1 and 3’ v2 chemistries (16.8 FPKM and 20.2 FPKM, respectively). The ddSEQ and Drop-seq methods had comparable dropout rates (25.0 FPKM and 26.7 FPKM respectively), while ICELL8 methods had the lowest sensitivity (37.9 FPKM/3’ DE and 112.1 FPKM/3’ DE UMI) (**Figure 3F and Table 1**). We observed similar trends across methods with the other three cell types, which had greater variance in read depth and transcript detection (**Supplemental Figure 4B-D**).

### mRNA Detection Affects the Fidelity of Single-Cell and Pseudo-Bulk Transcriptomes

We next investigated how well single-cell expression recapitulates immune signatures from bulk RNA-seq. For this purpose, we correlated expression of a set of marker genes (defined using bulk RNAseq data; see Methods) between bulk RNAseq and single cells. In general, cells with more genes detected had a better concordance to bulk RNA-seq immune signatures (**Supplemental Figure 5**). We observed higher Pearson correlation coefficients for 10x 3’ v3, 5’ v1 and ddSEQ methods against EL4, IVA12 and Jurkat bulk RNA-seq expression signatures (**Figure 4A**). ICELL8 3’ methods, with generally fewer genes detected, demonstrated the lowest correlation values. Overall, poorer correlation to TALL-104 bulk RNAseq was in line with fewer transcripts and genes detected for this cell type in the single-cell data.

**Figure 4:**
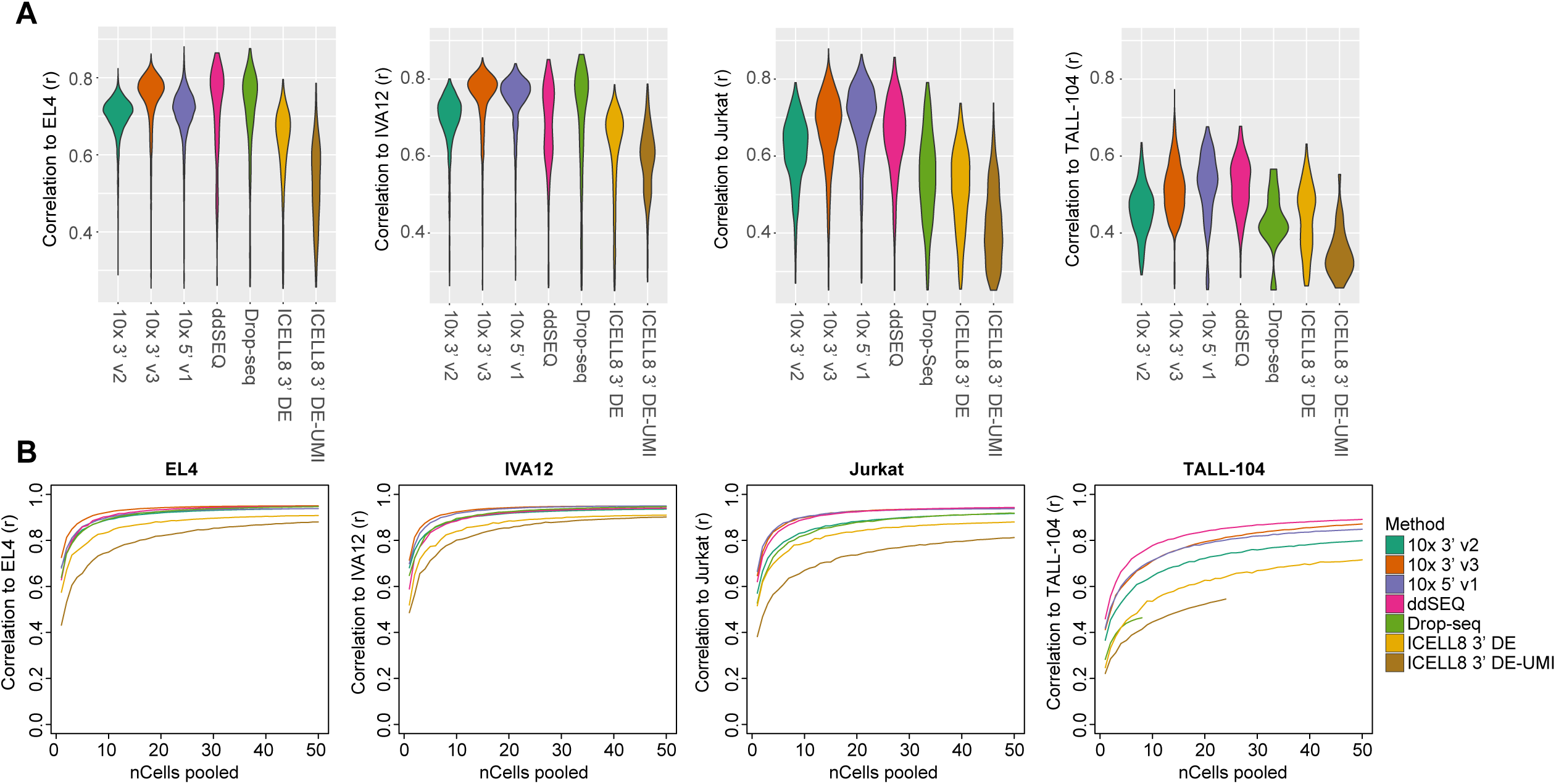
Correlation to bulk RNA-seq. (A) Pearson correlation (*r*) of cell identifiers (CIDs) to bulk RNA-seq data using highly-expressed variable genes. Only *r* values above 0.2 were included in plot. (B) Average Pearson correlation using all genes for aggregated data of 50 subsamples of up to 50 cells are plotted.

We further examined the correlation between pooled single-cell RNA-seq pseudo-bulk transcriptomes and bulk RNA-seq data using all genes. Averaging gene-expression profiles across single cells is commonly performed to compare data across experiments and is thought to resemble bulk data.

For EL4, IVA12 and Jurkat, most methods began to plateau around a correlation value of *r* = 0.9 with a pool of 10-20 cells (**Figure 4B)**. The maximum correlation values were lower for ICELL8 3’ DE (*r* = 0.90 and 3’ DE-UMI methods (*r* = 0.81-0.90) compared to other methods (r=0.92-0.95), and correlation was generally lower for TALL-104 cells in all methods, suggesting that lower mRNA detection sensitivity not only affects data fidelity at a per-cell level, but also impacts aggregated single-cell data. Although samples were prepared under identical conditions, we cannot rule out any effects of biological differences between samples. However, it is likely that higher variance in the detection of lowly expressed transcripts drives differences in expression observed in single-cell and bulk RNA-seq. Notably, our data indicates that this technical variance is not necessarily reduced by pooling across single cells and results from such analyses should be interpreted cautiously.

### Higher mRNA Detection Sensitivity Improves Identification of Differentially Expressed Genes

To assess the performance of differential expression analysis for each method, we focused on the two mouse cell types (EL4 and IVA12), because these cells had more similar sequencing depths compared to the two human cell types. We used the hurdle model proposed by Finak *et al*. [1] to identify differentially-expressed (DE) genes with an FDR < 10^−4^. For each DE analysis we sampled 199 cells, the lowest number of recovered cells by any method. Over 3,000 DE genes were identified in 10x Genomics methods, the highest among the methods tested, followed by Drop-seq (avg ∼2,700 genes) and ddSEQ (avg ∼2,800 genes), while the two ICELL8 methods had the fewest number of DE genes (avg ∼1,800 and ∼1,000 genes) (**Figure 5B and Table 1**). We observed similar trends with two alternative commonly-used tests for differential expression, a Mann-Whitney-Wilcoxon test [1] and a likelihood ratio test with an negative binomial generalized linear model [1, 2] (**Supplemental Figure 6B**).

**Figure 5:**
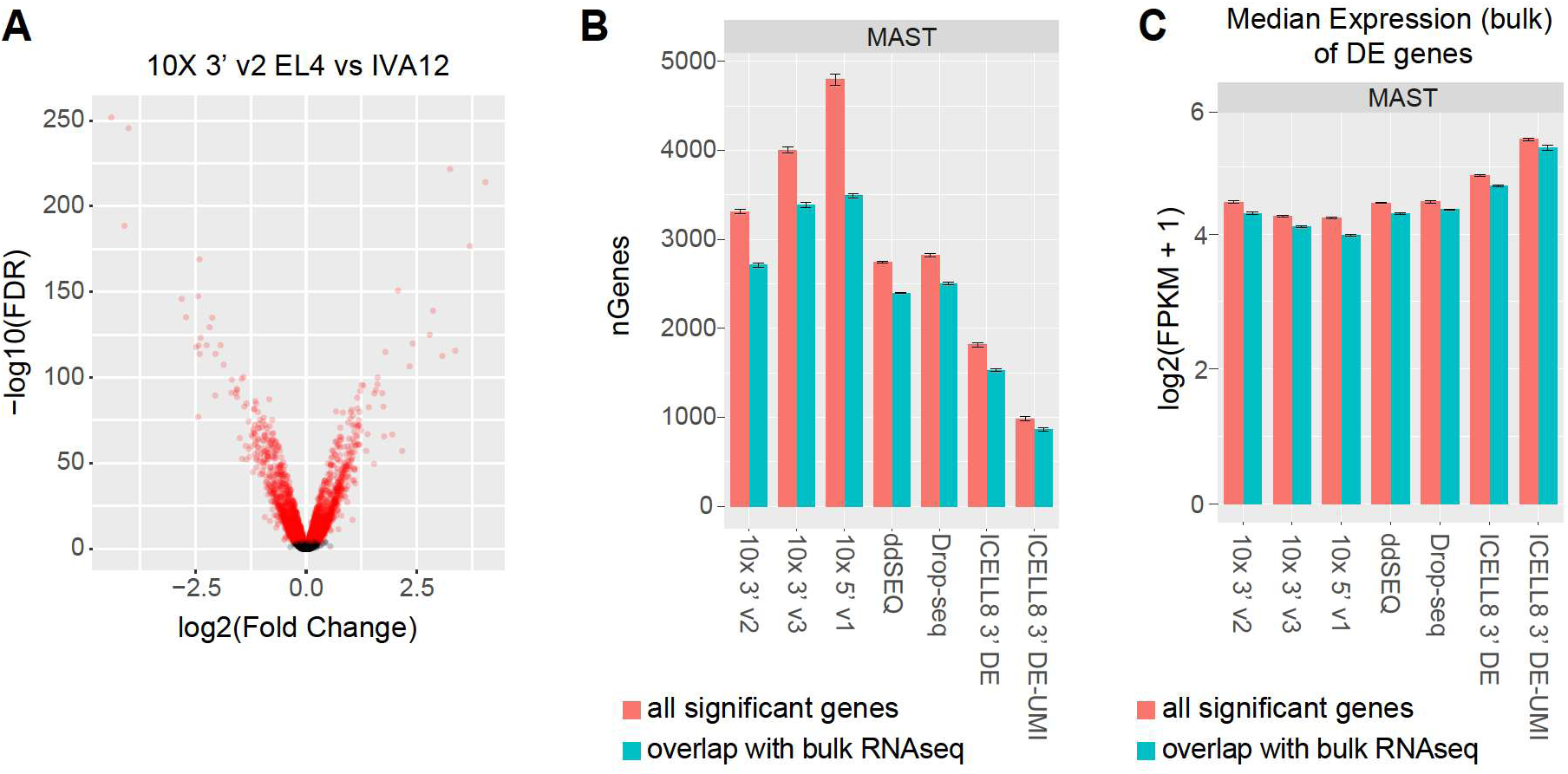
Differentially-expressed (DE) gene detection. (A) Fold change (FC) versus false discovery rate (FDR) calculated using a hurdle model (MAST) for mouse genes in EL4 vs IVA12 cells. Shown is a representative subsample of mouse cells (*n*=199) using the 10x 3’ v2 method demonstrating the criteria for declaring DE genes (FDR < 10^−4^); DE genes are highlighted in red. (B) Number of significant DE genes calculated using MAST between EL4 and IVA12 cells by method. Error bars represent the 95% confidence interval. The total number of significant DE genes are plotted in red, the number of DE genes with > 1.5 fold difference in expression in bulk RNA-seq (5,868 genes) are plotted in cyan. (C) Median bulk RNA-seq expression (FPKM) of all significant DE genes (red) or DE genes with > 1.5 fold difference (cyan). Error bars represent 95% confidence interval.

Performing DE analysis using all the cells obtained in each method increases the number of genes passing the significance threshold due to the increased statistical power (**Supplemental Figure 6D**). When we only consider 5,868 genes that have more than a 1.5-fold difference in bulk RNA-seq data, a proxy for ground-truth expression differences, the trend remains the same (**Figure 5D and Table 1**).

In general, we observed that fold-changes in single-cell data correlated well with gene expression differences in bulk RNA-seq data, especially for genes with higher expression levels (**Supplemental Figure 6A**). In contrast, genes with low expression correlated poorly with smaller fold changes observed in the single-cell data, consistent with higher dropout probabilities for lowly-expressed transcripts. Also, the distribution of FPKM values is generally higher for DE genes from single-cell data compared to genes with at least 1.5 fold-change in bulk RNA-seq (**Supplemental Figure 6E**), indicating that all methods exhibit a considerable type II error rate. Furthermore, we find the lowest median FPKM in bulk RNA-seq for DE genes from the methods with the highest detection sensitivity (10x 3’ v3 (median = 3.43 FPKM) and 10x 5’ v1 (median = 3.45 FPKM), and the highest median FPKM for the ICELL8 3’ DE-UMI method (median = 4.91 FPKM), which had the lowest transcript detection sensitivity (**Figure 5C and Supplemental Figure 6C**).

### mRNA Detection Sensitivity Varies Across Heterogenous Cell Types

Many immune single-cell experiments profile an undefined mixture of cell types that potentially vary in mRNA content. Efficient cell recovery across diverse cell types is important to accurately characterize the diversity of these populations. We next compared the differences in cell recovery between the four cell types included in our sample mixture. In particular, TALL-104 cells are smaller (5µm diameter) than the other three cell-types (EL4/IVA12 - 11µm, Jurkat - 10µm diameter) and, in our hands, more difficult to culture, with viability rates under 80% and slow growth. Across all experiments, TALL-104 cells had the lowest distribution of reads, UMIs, and genes recovered (**Supplemental Figure 1C and 4A**), such that they were more susceptible to exclusion based on read or UMI thresholding of CIDs to distinguish cells from ambient noise.

We classified cells by correlating their expression profile to gene signatures from bulk RNA-seq. This enabled us to examine the recovery of each cell type with common thresholding points on the log-log curve of total reads or UMI vs rank ordered CIDs. In droplet-based methods, thresholding removes a large fraction of CIDs that are derived from droplets containing a barcoded bead, but no cell. Two points are commonly used as thresholds: the knee point where the signed curvature is minimized and the inflection point where first derivative is minimized [1] within a given range of total reads or UMIs (**Figure 6A and Figure S2A**). While the fraction of classifiable cells on each side of these two thresholds varied across experiments, both thresholds were able to capture EL4, IVA12 and Jurkat cells (**Figure 6B**). Notably, most TALL-104 cells would be removed by thresholding using the stringent knee point, with only one experiment having any TALL-104 cells above this threshold. While the more permissive inflection point performed better at capturing TALL-104 cells, all TALL-104 cells would be considered ambient noise using this threshold in samples (**Figure 6B**).

**Figure 6:**
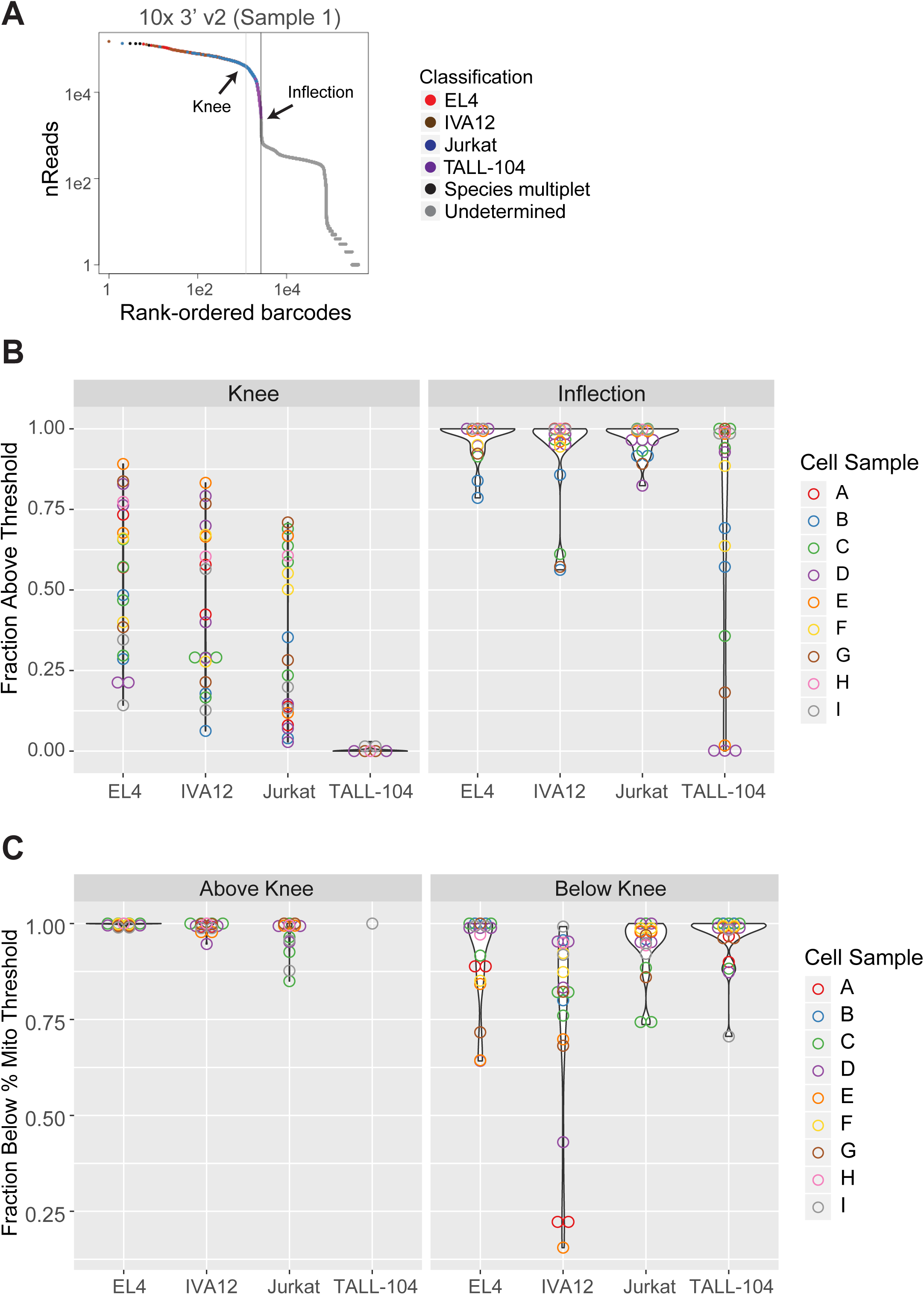
Cell recovery by cell identifier (CID) thresholding. (A) Example of using the transposed log-log empirical cumulative density plot of the total counts of each CID to identify cell-containing droplets. Common thresholding points, the ‘knee’ and the ‘inflection’ are indicated with arrows. The knee is the point at which the signed curvature is minimized, the inflection is the point at which the first derivative is minimized. (B) The fraction of cells above the knee or inflection are plotted. (C) Fraction of cells below mitochondrial rate threshold (listed in **Supplemental Table 4**) relative to knee point. Samples are colored by cell sample mixture listed **in Supplemental Table 2**.

As low mRNA recovery might reflect poor cell health, e.g., due to mechanical stress in cell preparation, we also examined the fraction of cells below each threshold that had a high rate of mitochondrially encoded UMIs or reads, an indication of broken or poor-quality cells (**Figure 6C**). Accordingly, a large fraction of cells removed by the knee point cutoff had a high mitochondrial rate. However, significant numbers of cells had reasonable mitochondrial rates, including many TALL-104 cells. This suggests that lower transcript recovery in TALL-104 cells is related, at least in part, to lower overall mRNA content and not cellular damage, although we cannot completely rule out other cell-quality issues that do not affect the mitochondrial rate. Overall, some cell populations could be lost when thresholding based on total UMI or read count is too stringent. It would be beneficial to include more cells at the initial CID selection step and filter cells more stringently in downstream analyses with other cell-quality criteria to avoid loss of cell populations with low mRNA content. Of note, this issue does not affect ICELL8 methods as all cell-related barcodes are known *a priori* when cell-containing wells are selected for processing.

In heterogeneous populations, mRNA capture rates and read depths may vary across subpopulations. We explored differences in mRNA detection sensitivity across the four cell types in our samples. As it is common in single-cell profiling of mixed populations, we observed differences in read and UMI recovery across cell types in each method (**Figure 7A**). When the entire data is used to model dropout rates, we find that cell types with the lowest read distributions, such as TALL-104 cells, have increased dropout probabilities and GD_50_ levels across all seven methods tested (**Figure 7B and Supplemental Figures 4B & 4D**). We hypothesized that differences in dropout rates were predominantly driven by differences in mRNA detection rates and compared cells from each cell type with similar numbers of UMIs. Cells were separated into six quantile bins based on the number of UMIs (**Figure 7C**) and dropout rates for each cell type were modeled. Because there were few TALL-104 cells recovered in many samples (**Supplemental Table 4**), we focused on data from the 10x 3’ v3 method which had sufficient numbers of cells available for analysis. We found that with increasing number of total UMIs, GD_50_ values and dropout rates decreased. Notably, GD_50_ levels were similar across cell-types within a bin (**Figure 7D**). Slight differences in GD_50_ are related to variation in mean number of UMIs for a particular cell type. TALL-104 cells, which fell into the two lowest bins due to the low numbers of transcripts detected, had similar dropout rates as other cell types in the same bin (**Figure 7D & 7E**).

**Figure 7:**
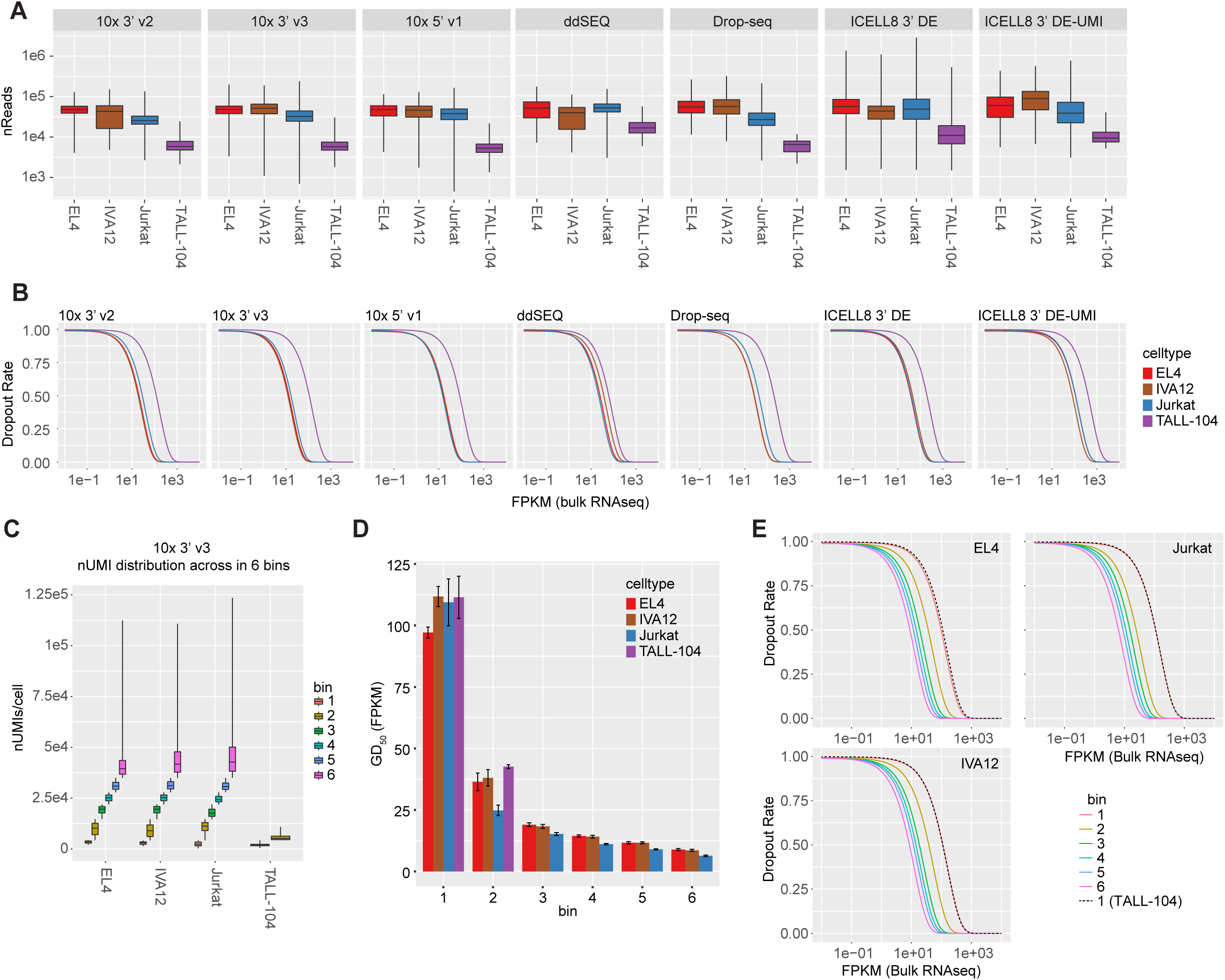
Dropout rates by cell type. (A) Distribution of reads across cell types is plotted by method. (B) Dropout rate models for cell types are shown. (C) 10x 3’ v3 cells were binned by number of unique molecular identifiers (UMI) and distributions of nUMIs for each cell type in each bin are plotted. (D) Gene Detection 50 (GD_50_) rates, expression level at 0.5 probability of the dropout model, are plotted for each cell type in 10x 3’ v3 experiments by bin. (E) Dropout models in each bin for EL4, IVA12 and Jurkat cells are plotted along with the model for TALL-104 cells in bin 1.

## Discussion

In this paper we explored several important quality metrics of single-cell RNA-seq methods: efficiency of cell recovery, library efficiency and mRNA detection sensitivity. High recovery of cells put into a system and minimal loss of reads due to noise are important, especially to limited samples with few cells. The differences in performance we observed across these methods are directly related to the design of cell and mRNA capture. To partition cells, the methods we tested either use microfluidics to generate nanoliter sized droplets or to partition cells on microwell chips. Ideally, those microreactors contain exactly one bead and one cell. In practice, however, the number of cells per microreactor approximately follows a Poisson distribution. While the loading probability of cells is similar across these methods, the distribution of barcoding oligonucleotides varies. The loading statistics of Drop-seq and ddSEQ follow a Poisson distribution, while 10x Chromium chips load beads in a sub-Poissonian fashion. The latter enables an increased theoretical capture rate of ∼60%. Sparser loading of barcoding beads in ddSEQ and Drop-seq minimizes the occurrence of bead doublets, but at the expense of lower maximum recovery rates of ∼3% and ∼5% respectively. Oligonucleotide loading is tightly controlled on ICELL8 chips with the pre-printing of oligonucleotides, providing *a priori* knowledge of cell-related CIDs when coupled with cell imaging. The ability of the ICELL8 to selectively process a subset of wells, those containing cells identified by fluorescence imaging, greatly improves this method’s library efficiency compared to techniques that process all partitions. Accordingly, we observed the highest fraction of cell-related reads in ICELL8 libraries, especially compared to ddSEQ and Drop-seq methods with a large fraction of bead containing droplets lacking a cell and increased potential for ambient RNA. Quality of single-cell suspensions are also important factors to these metrics. Variable cell viability and inefficient cell quantification of our samples may negatively impact cell-capture and multiplet rates and explain the discrepancy between expected and observed rates.

In our experiments, we observed the highest mRNA-detection sensitivity in the 10x 3’ v3 and 5’ v1 methods, with the highest numbers of transcripts and genes detected and lower probabilities of gene dropouts at lower expression levels. Our results corroborate previous reports about the performance of some of the methods assessed in our work [1-2]. Here, we extend these findings by demonstrating an increased sensitivity of the more recent 5’ v1 and 3’ v3 methods, which also validates claims made by 10x Genomics. Further, we found ICELL8 methods have the lowest mRNA detection sensitivity of the methods tested for the assayed immune cell types. Of note, this is partly in disagreement with two papers reporting better performance of the ICELL8 3’ DE method relative to 10x 3’ v2 and Drop-seq [1, 2]. Differences to the performance we observed may be related to cell types used in each study. For example, ICELL8 3’ DE detected significantly fewer genes per cell compared to 10x 3’ v2, ddSEQ and Drop-seq in B-cells in Mereu *et al*.[1], which is on par with our findings.

Gene detection rates may be increased by greater sequencing depths, particularly for low expressed genes (**Supplemental Figure 2B and Supplemental Table 3**). However, high-throughput methods aim to sequence many cells concurrently for a broad exploration of populations, at the expense of the completeness of individual transcriptional profiles. Here, libraries are not routinely sequenced to full saturation due to high sequencing costs. To be able properly assess mRNA detection sensitivity, we normalized samples to a common sequencing depth of ∼50,000 reads per cell by downsampling raw reads. Additional iterations of this stochastic process showed little variation in the resulting analysis (**Supplemental Table 5**), suggesting our normalization step did not introduce any technical bias. Notably, the resulting sequencing depth is typical for common high-throughput single-cell RNA-seq experiments. Therefore, our data can provide expectations for mRNA and gene detection rates in experiments with a similar sequencing depth using other immune cells.

Multiple aspects of single-cell RNA-seq protocols such as efficiencies in mRNA capture, reverse transcription, and cDNA amplification can affect the overall mRNA detection sensitivity. Efficient mRNA capture may be impacted by the template switch mechanism, as only first strand cDNAs which have successfully switched templates can be amplified. ddSEQ, the sole method we tested that does not utilize template switching, is not as sensitive as the 10x Genomics methods, possibly due to other technical differences. Another source of inefficiency may arise in the reverse transcription cleanup step prior to cDNA amplification. We found that the addition of a primer digestion step to the ICELL8 3’ DE protocol in the 3’ DE-UMI method decreased the mRNA detection sensitivity. Additional improvements to mRNA capture such as improving oligonucleotide chemistry for mRNA capture and cDNA amplification may enhance mRNA detection sensitivity and improve single-cell RNA-seq techniques in the future.

Increasing the sensitivity of mRNA detection greatly benefits downstream analyses of immune profiling datasets. Sampling transcriptomes with high fidelity results in a greater likelihood of detecting rare transcripts for identifying DE genes at lower expression levels. In general, our results show that expression profiles of cells with high mRNA content generated by methods with a high mRNA detection rate, correlated well to bulk-RNA-seq data. Also, the number of DE genes as well as the overall correlation in fold-change differences to bulk RNA-seq improved with higher mRNA detection sensitivity. Here, all 10x Genomics methods, which had the highest mRNA detection sensitivity, exhibited a high correlation to bulk RNA-seq data as well as more DE genes with a lower range of expression levels in bulk data. Notably, our results revealed that the higher variance in the detection of lowly expressed transcripts commonly observed in techniques with lower sensitivity is not necessarily overcome by pooling across single cells when performing pseudo-bulk analyses. Strengthening the underlying mRNA detection sensitivity can improve downstream analyses to identify marker genes as well as classify subtle immune subtypes and cell states with small, but significant differences in gene expression, and can facilitate the identification of novel immune subpopulations [1].

Importantly, our data also provides insight into the performance of single-cell techniques across heterogenous populations of immune cells. Although the immune cell lines used in this study may differ from lymphocytes found *in-vivo*, the standardized cell culture conditions for these cells helps reduce expression variability compared to primary cells and facilitated data analysis. Nonetheless, our results provide better guidance for immune profiling in contrast to the higher mRNA content cell lines such as carcinoma or stem cells commonly used in previous comparison papers [8, 11, 12]. The inclusion of small TALL-104 cells allowed us to assess the sensitivity of these methods in subpopulations with comparatively low mRNA levels. We observed that relaxing CID filtering criteria based on total UMI or total read counts can improve recovery of TALL-104 cells for downstream analyses. Notably, smaller immune cells such as TALL-104 cells have a higher gene dropout rate that is related to the number of transcripts captured from a cell. Thus, additional quality metrics also need to be calibrated carefully for the identification of small immune cell types. Cell imaging on the ICELL8 cx to identify otherwise challenging cells of interest such as TALL-104, can also be used to recover populations of small cells. In general, TALL-104 cells exhibited lower mRNA detection rates and higher dropout rates. Thus, we can expect that other immune cell types with low mRNA content exhibit similar dropout rates as other immune cells with a comparable rate of transcript detection.

## Conclusions

Our comparison of data from seven high-throughput single-cell methods can help guide method selection for immune profiling experiments. Our data can provide reasonable predictions of transcript and gene detection rates for immune cells, as well as insight into performance across heterogenous lymphocyte populations with varying mRNA content. Our results suggest looser thresholding of CIDs in droplet-based methods can be beneficial to retain cell populations with low-mRNA content. Additionally, smaller cells such as TALL-104 cells have a higher gene-dropout rate that is related to the number of transcripts captured from a cell.

Each method we tested showed advantages that could benefit immune cell profiling. In this study, 10x Genomics methods had the highest cell recovery and mRNA detection sensitivity, making these techniques particularly suited to experiments with limited samples and experiments that require detection of genes with lower expression levels. Here, the performance was comparable between the 10x 5’ v1 chemistry and 3’ v3 methods, making the 5’ v1 chemistry an appropriate substitute when pairing gene expression analysis with TCR/BCR clonotyping. 10x Genomics and Illumina/Bio-Rad (ddSEQ) sell reagents in kits, facilitating adoption of these methods, but limiting customization of protocols.

Takara Bio also sells reagent kits for the ICELL8, however, protocols on the instrument are customizable allowing for greater flexibility. Drop-seq is also an open system that is fully customizable and custom reagents such as target-capture oligonucleotide beads [1, 2] can be easily integrated into the protocol. The fluorescent imaging capabilities of the ICELL8 cx enable the pairing of sequencing and imaging data, in downstream analysis. Our ICELL8 experiments demonstrated high library efficiencies, with a large fraction of reads assignable to cells and potential utility to recover low-mRNA-content cells, such as TALL-104 cells, that are more susceptible to stringent read and UMI thresholding. Overall, our data shows that all methods exhibit specific strengths which can be aligned with experimental goals, sample limitations, and budgetary constraints.

## Materials and Methods

### Cell Culture

All cell lines were acquired from ATCC. EL4 (ATCC TIB-39) cells were cultured in RPMI-1640 + 2 mM L-glutamine + 10% FBS + 1.7 ul 2-mercaptoethanol per 500ml media. IVA12 (ATCC HB-145) cells were cultured in DMEM + 10% FBS + 1X P/S. Jurkat (ATCC TIB-152) cells were cultured in RPMI-1640 + 10% FBS + 1X P/S. TALL-104 (ATCC CRL-11386) cells were cultured in IMDM + 15% FBS + 1x L-Glu + 200U/ml IL-2. Prior to processing cells, cells were washed in 1X PBS and cell concentration and viability were determined using a Countess (Invitrogen). Cells were mixed at a 1:1:1:1 ratio based on viable counts and resuspended in PBS + BSA solution according to manufacturer’s guidelines.

### Chromium

Cells were resuspended in PBS with 0.04% BSA at a stock concentration within the recommended range (typically ∼1e6 cells/mL) and loaded at a volume to target between 2000-6000 cells depending on sample. Libraries were prepared according to manufacturer’s instructions for each chemistry. Libraries were sequenced on a NextSeq500 or NovaSeq (Illumina) according to manufacturer’s guidelines: 3’ v3 - 28×8×0×91, 3’ v2 - 26×8×0×98, and 5’ v1 - 26×8×0×110.

### ddSEQ

Cells were resuspended in PBS + 0.1% BSA and loaded at a concentration of either 2000 or 2500 cells/ul. Libraries were prepared according to manufacturer’s instructions. Libraries were sequenced on an Illumina NextSeq500 (68×8×0×150) at a 3pM concentration with provided custom read 1 primers.

### Drop-seq

Libraries were prepared following the McCarroll Lab Drop-seq protocol (http://mccarrolllab.org/Drop-seq/) [1], with cells and beads encapsulated using the Dolomite scRNA-Seq system. Oligo beads (ChemGenes) contained the original Drop-seq polyT primer with a VN anchor at the 3’ end (TTTTTTTAAGCAGTGGTATCAACGCAGAGTACJJJJJJJJJJJJNNNNNNNNVTTTTTTTTTTTTTTTTTTTTTTTTTTTTTT VN). Cells were resuspended in PBS + 0.01% BSA and loaded at a concentration of either 150 or 300 cells/ul. For encapsulation, cell and bead solutions were loaded at 30 ul/min and encapsulation oil was loaded at 200 ul/min. Libraries were sequenced on a NextSeq 500 (Illumina) (21×8×0×138) at a 3pM concentration with custom Read1 Drop-seq primers (GCCTGTCCGCGGAAGCAGTGGTATCAACGCAGAGTAC).

### ICELL8 CX

Cells were resuspended in 1X PBS and loaded at a final concentration of 2,800 cells/ml. Only wells containing single cells as determined by the Cell-Select software using default settings were processed. Libraries were prepared using the Takara Bio SMARTER ICELL8 cx 3’ DE user manual or an alternate protocol that separates the initial reverse transcription reaction from cDNA amplification. In short, after RT, cDNA was removed from the chip and cleaned and concentrated with a Zymo Clean & Concentrator-5 kit. cDNA was then treated with 20U of Exonuclease for 30 min at 37°C and the enzyme was deactivated with 20 min at 80°C. cDNA was then amplified and tagmented using the Illumina Nextera XT kit for the final sequencing libraries. Libraries were sequenced at 25×8×0×131.

### Bulk RNA sequencing

RNA was isolated from cells using the RNeasy kit (Qiagen). Libraries were generated using 1ug of total RNA using a modified Illumina TruSeq Stranded mRNA protocol. Reverse transcription was performed with the addition of RNaseOut (Invitrogen) and actinomycin-D (MP Biomedicals). The resulting product was cleaned using AMPure RNAClean beads. Additionally, second-strand synthesis was performed using dUTP instead of dTTP and an additional USER (New England Biolabs) digestion step was incorporated after size-selection. Libraries were sequenced at 101×6×0×101 on a HiSeq (Illumina) to a minimum depth of 30 million reads per sample.

### Read alignment and transcript counting

All sequencing data was aligned to a combined human/mouse reference genome obtained from 10x Genomics: reference “cellRanger_1.2.0” composed of hg19 with Ensembl 82 and mm10 with Ensembl 84. Bulk data was aligned using STAR v2.5.1b and quantified using featureCounts v1.6.3 [1].

For normalizing single-cell libraries, we considered the fact that cell types with low mRNA content are more prone to drop-outs and thus, may compromise proper normalization based on the mean read count per CID. Thus, robust library scaling factors were derived by using only cells with sufficiently high mRNA content. For this purpose, we calculated Gaussian kernel density estimates with a smoothing bandwidth determined via the Normal Reference Distribution method as provided by R v4.0.2. Local maxima of the density function were sorted in decreasing order. The first significant mode, *d*_i_, was considered the sequencing depth of the cell population with the highest mRNA content for sample *i* (**Figure S1A**). Scaling factors were then derived by *s*_i_ = min_i_(*d*_i_)/*d*_i_. FASTQ files were downsampled by factor *s*_i_ using seqtk v1.3-r106. This resulted in libraries with around 50,000 reads per cell. Downsampling was repeated three times for four representative samples for an assement of the margin of error. Alignment statistics for normalized data were generated using PicardTools CollectRNA-seqMetrics using the aligned (and filtered) BAM files from each pipeline.

Downsampled FASTQ files were further processed using method-specific pipelines with parameters set as recommended. All pipelines employ STAR [1] for the alignment step, but are tailored to identify method-specific barcodes and count transcripts. Chromium data was processed using Cellranger v3.0.2 (with STAR v2.5.1b); ddSEQ, Drop-seq and ICELL8 data were processed using the Drop-seq_Tools pipeline v2.3.0 (with PicardTools v2.18.14 and STAR v2.4.2a). ddSEQ CIDs and UMIs were extracted using ddSeeker v0.9.0 [1]. ICELL8 read and UMI count matrices was generated using mappa v0.9 software (with STAR v2.7.0b).

To assess step-wise quality metrics of each method’s original data we applied a uniform pipeline (**Supplemental Table 3**). First, we demultiplexed and aligned sequenced reads using STAR as recommended by each scRNA-seq method. Here, ddSEQ data was processed using SureCell RNA Single-Cell v1.1.0 (with STAR v2.5.2b). Then, high-quality reads were filtered by MAPQ scores (MAPQ = 255 for 10x, Drop-Seq, and ICELL8; MAPQ = 50 for ddSEQ) using samtools 1.9 [1] and mapped against mouse (Ensembl 84) and human (Ensembl 82) gene annotations using featureCounts from the subreadpackage 2.0.1 with parameters “-t exon -g gene_id -C -p --primary”. We wrote custom Java code (v JDK 11.0.1) to generate read and UMI count matrices. Here, we considered potential sequencing errors in UMIs and corrected these as follows: reads got grouped by <barcode, gene, UMI> tuples. If two groups had the same <barcode, gene> pair, but their UMIs differed by a single base, the UMI of the smaller group was corrected to the UMI of the larger group.

### Cell classification

Cells were assigned to one of four input cell classes by their similarity to cell type signatures from bulk RNA-seq data. First, we selected highly expressed genes with FPKM > 50 in any bulk RNA-seq sample. Next, gene expression was contrasted between bulk RNA-seq samples from the same species (EL4 vs IVA12 and Jurkat vs TALL-104) and we filtered 184 highly variable genes (93 mouse, 91 human) with a ln fold difference > 3 between the two cell types. Pearson correlation, *r*, was calculated between each gene expression vector of each cell **x**_i_ ∈ ℝ ^8^ and each gene expression vector of each bulk RNA-seq sample **y** ∈ ℝ ^8^ : *f*_i_ **y** = *r*(ln (**x**_i_ + 1), ln (**y** + 1)). A cell type was assigned using the following four classification rules derived from the correlation coefficient distributions:

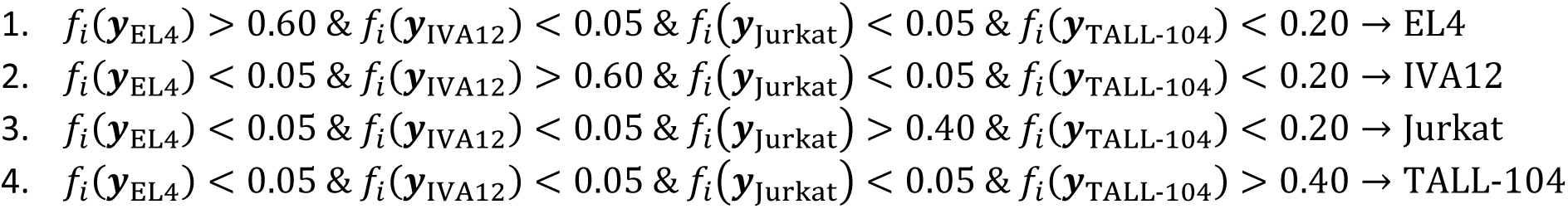

We relaxed these rules for ICELL8 data to account for overall lower CID numbers and method-specific distribution differences:

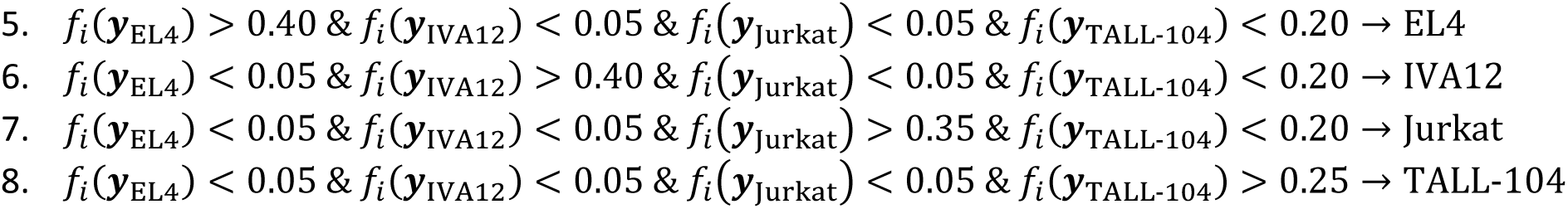

Cells with two or more assigned cell types, were removed. For each sample, we classified the top *n* CIDs ranked by total read count with *n* = 2 × number of expected cells.

To analyze the thresholding method using the transposed log-log empirical cumulative density plot of the total read counts of each CID, we calculated the knee and inflection points as described in Lun et al. [1]. Briefly, the knee and inflection points are interpreted as determinants of the range in which the curve transitions between cell-containing droplets/wells with high mRNA content and empty droplets/wells with ambient RNA. The total count per CID is modeled as a function of decreasing CID rank by fitting cubic smooth splines with 20 degrees of freedom. The knee point is defined as the point on the curve where the signed curvature is minimized, the inflection point is defined as the point where the first derivative of the spline basis functions is minimized. We defined a lower bound for fitting the smooth splines as the minimum number of total reads of all classified CIDs. Calculations were performed using the R package DropletUtils v1.6.1. For the analysis of the original (i.e., not down-sampled) data described in **Supplemental Figure 2**, CIDs above the inflection point were considered genuine cells.

### Doublet rate estimation

CIDs were classified as multi-species multiplets if the number of total counts from each species exceeded the 10^th^ percentile of the distribution for that species. The total count distributions were calculated using cells assigned in the step described above. Multiplet rates were calculated by taking CIDs above the inflection point and dividing the number of multi-species multiplets by the total number of cells. To obtain the true multiplet rate that considers non-detectable intraspecies multiplets and accounts for differing proportions of human and mouse cells, this fraction was divided by an adjustment factor *λ*:

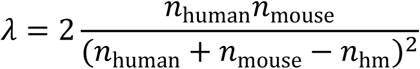

where *n*_human_ is the number of human cells, *n*_mouse_ is the number of mouse cells, and *n*_hm_ is the number of inter-species multiplets.

### Cell recovery rates

Cell recovery rates were calculated as the number of CIDs above the inflection point divided by the total number of cells loaded onto a system. Theoretical capture rates for 10x Genomics and ddSEQ methods are based on expected recovery numbers given in the user manual. Theoretical capture rates for DropSeq were based on a 5% droplet occupancy of oligo beads. Theoretical capture rates for iCELL8 protocols were calculated based on a Poisson distribution of wells with single cells based on an average cell occupancy of 1 cell per well.

### Pseudo-bulk analyses

Pseudo-bulk analyses analyzing correlation to bulk RNA-seq data and gene detection rates were performed by summing UMI counts across multiple cells. A subsample with at least 100 classified cells or the maximum number of classified cells recovered that meet the mitochondrial rate threshold were selected from each method. Mitochondrial rate thresholds were determined on a per-sample basis based on distribution of rates (**Supplemental Table 4**). Various numbers of cells (1-50) were randomly sampled from this pool and expression values were averaged. The aggregated expression matrix was used for analyzing its correlation to bulk RNA-seq and for quantifying the number of detected genes. Mean values across 50 iterations for these metrics were used for visualization.

### Dropout modeling

Dropout rates denote the fraction of missing values in a gene’s expression vector. We estimated the dropout rate for each gene from the species of the cell type considered. Cells included in the analysis were filtered by fraction of mitochondrial counts to remove poor quality cells. Dropout rate of bulk RNA-seq data was modeled by fitting the function *f*(*x*) = *a* ∗ exp(–*b* ∗ *x*) where *x* is the expression level using nonlinear least squares. GD_50_ FPKM numbers were calculated as 0.5 = *a* ∗ exp(–*b* ∗ *x*) using the fitted coefficients *a* and *b*. Dropout rates were similarly calculated for single-cell RNA-seq data by binning cells by mRNA detection rates. 10x 3’ v3 cells were placed into six bins based on distribution percentiles resulting equivalent numbers of cells in each bin. Dropout models were calculated for a random subset of 50 cells for each cell type with at least 50 cells in each bin and results were averaged across 50 iterations.

### Differentially expressed gene identification

Statistical differences between gene expression of EL4 and IVA12 cells were identified using the hurdle model provided in the MAST R package v1.12.0 [1], a Wilcoxon rank-sum test, or the negative binomial generalized linear model available in the MASS R package v7.3-51.5. Genes that had an FDR-adjusted *p*-value < 1e-4 were declared differentially expressed. Cells from multiple replicates from each method were pooled in order to maximize sample sizes. Downsampling of cells was performed to the smallest number of observed cells from a single cell type (*n* = 199); this step was repeated 10 times to assess the error margin. UMI count data was used for 10x, ddSEQ, Drop-seq, and ICELL8 3’ DE-UMI samples, read count data was used for ICELL8 3’ DE samples. Expression count matrices were normalized by library size factors (i.e., total counts per cell), multiplied by 10^4^, and log-transformed by log (*x* + 1). Log-normalized count matrices were subjected to MAST, normalized count matrices were used for the Wilcoxon-rank sum test, and raw count data was input to the negative binomial generalized linear model.

## List of abbreviations

RNA-seq: RNA sequencing
PBMC: Peripheral blood mononuclear cell
DE: Differentially-expressed
UMI: Unique Molecular Identifier
CID: Cell identifier
cDNA: Complementary DNA
mRNA: Messenger RNA
PCR: Polymerization chain reaction
UTR: Untranslated region
**GD**_**50**_: Gene detection 50
FPKM: Fragments per kilobase of transcript per million mapped reads
TCR: T-cell receptor
BCR: B-cell receptor

## Declarations

### Ethics approval and consent to participate

Based on Amgen guidelines for ethics approval and consent to participate, the need for approval was waived in this study because no clinical samples were included.

### Consent for publication

All authors read and approved the final draft of the manuscript. This manuscript also has been reviewed and approved by Amgen Final Publication Review (FPR) process.

### Availability of data and materials

All data generated or analyzed during this study are included in this published article and its supplementary information files

### Author Contributions

TMY, DL, and CML designed the experiments and drafted the manuscript. TMY, DL, DCE, and CML performed data analysis. HZ, DB, and PM carried out the NGS run with TMY and DL and coauthored the section of methods for the manuscript. DCE, OKY and OH set up the bioinformatics analysis pipeline and provided input and assistance in preparation of this manuscript. SW and CML initiated the study and made contributions to experiment design and advised on data analysis, validation, and manuscript preparation.

## Acknowledgments

The authors would like to thank Wenny Chou for assistance in data generation and Ian Driver and Wenjun Ouyang for the valuable inputs to experimental design and manuscript preparation.

## Figure Legends

**Supplemental Figure 1:**
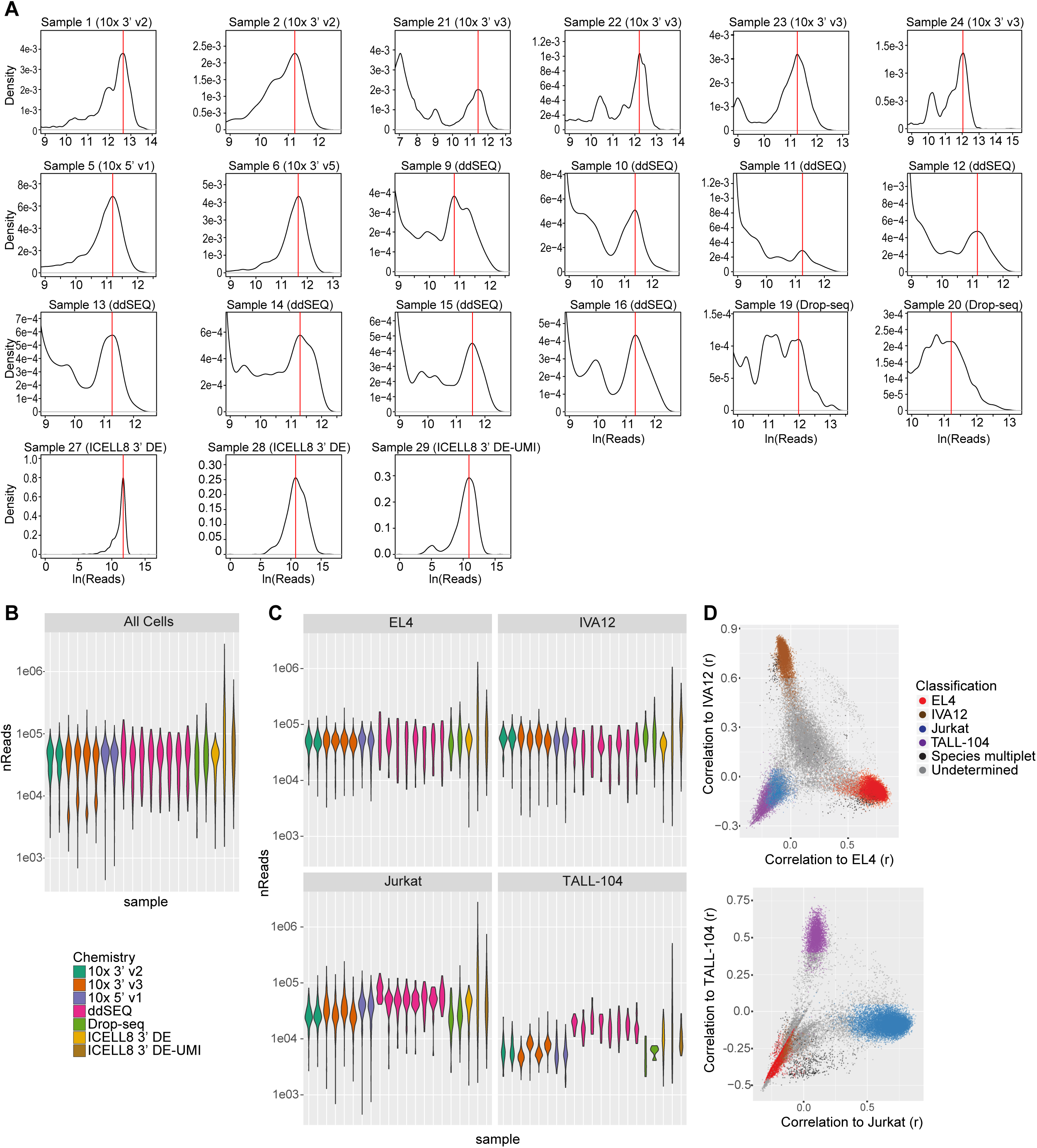
Read-depth normalization and cell classification. (A) Determined read distribution peaks used for read-depth normalization. (B) Normalized read distribution per experiment for classified cells. (C) Normalized read distributions per experiment and cell type. (D) Correlation of gene counts to bulk-RNA-seq data for all experiments. Top cell identifiers by numbers of reads (2x number of expected cells) were included in the plot.

**Supplemental Figure 2:**
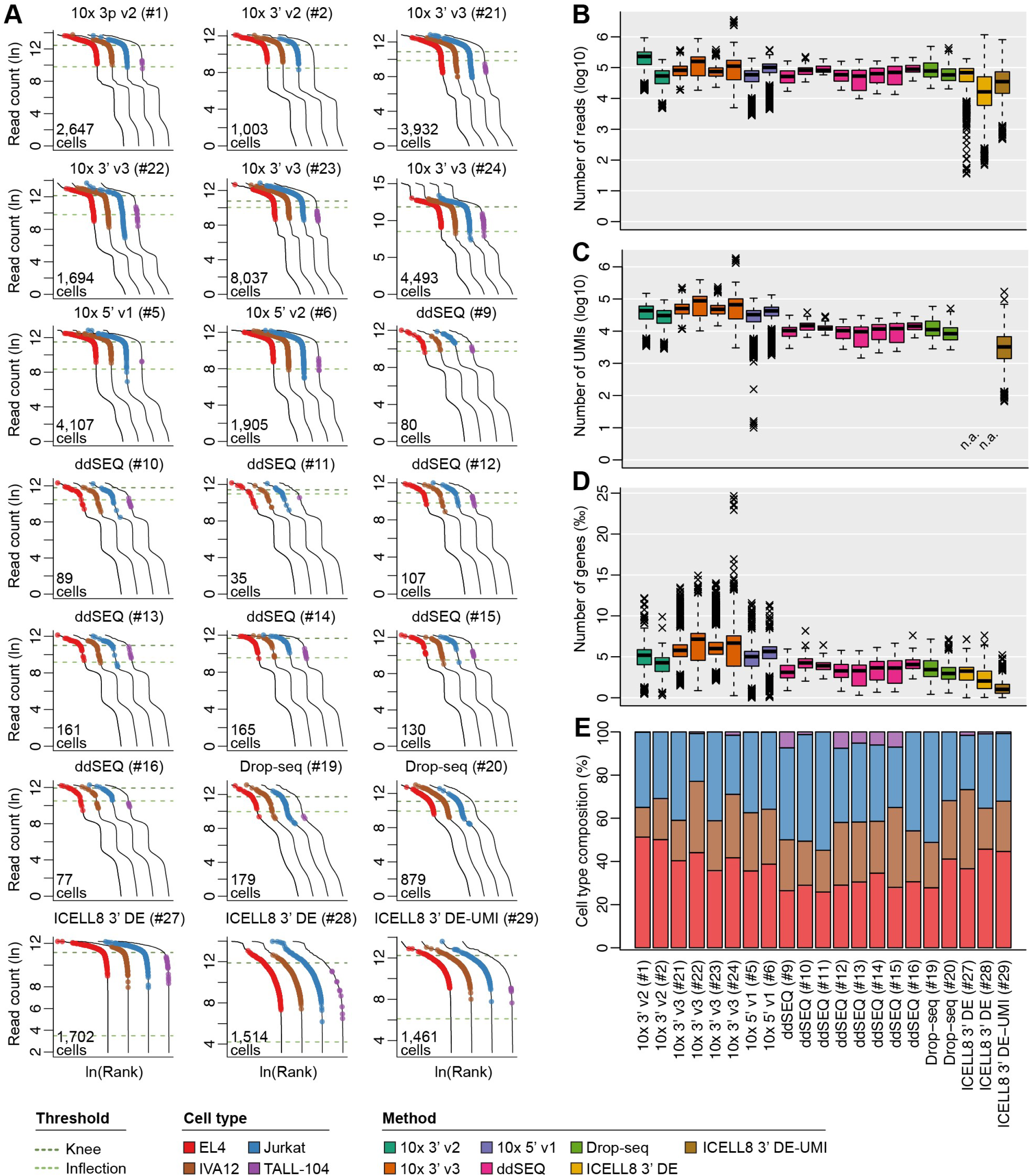
Cell metrics of pre-normalized data. **(A)** Transposed log-log empirical cumulative density plot of the total counts of each CID. The curve is shifted in each plot to highlight each cell and cell type that was determined by correlation to bulk RNA-seq using correlation thresholds determined in downsampled data. Knee and inflection points are indicated. (B-D) Boxplots show number of reads, number of UMIs, and number of detected genes per cell without normalization of library read depth. (E) Cell type composition of each sample.

**Supplemental Figure 3:**
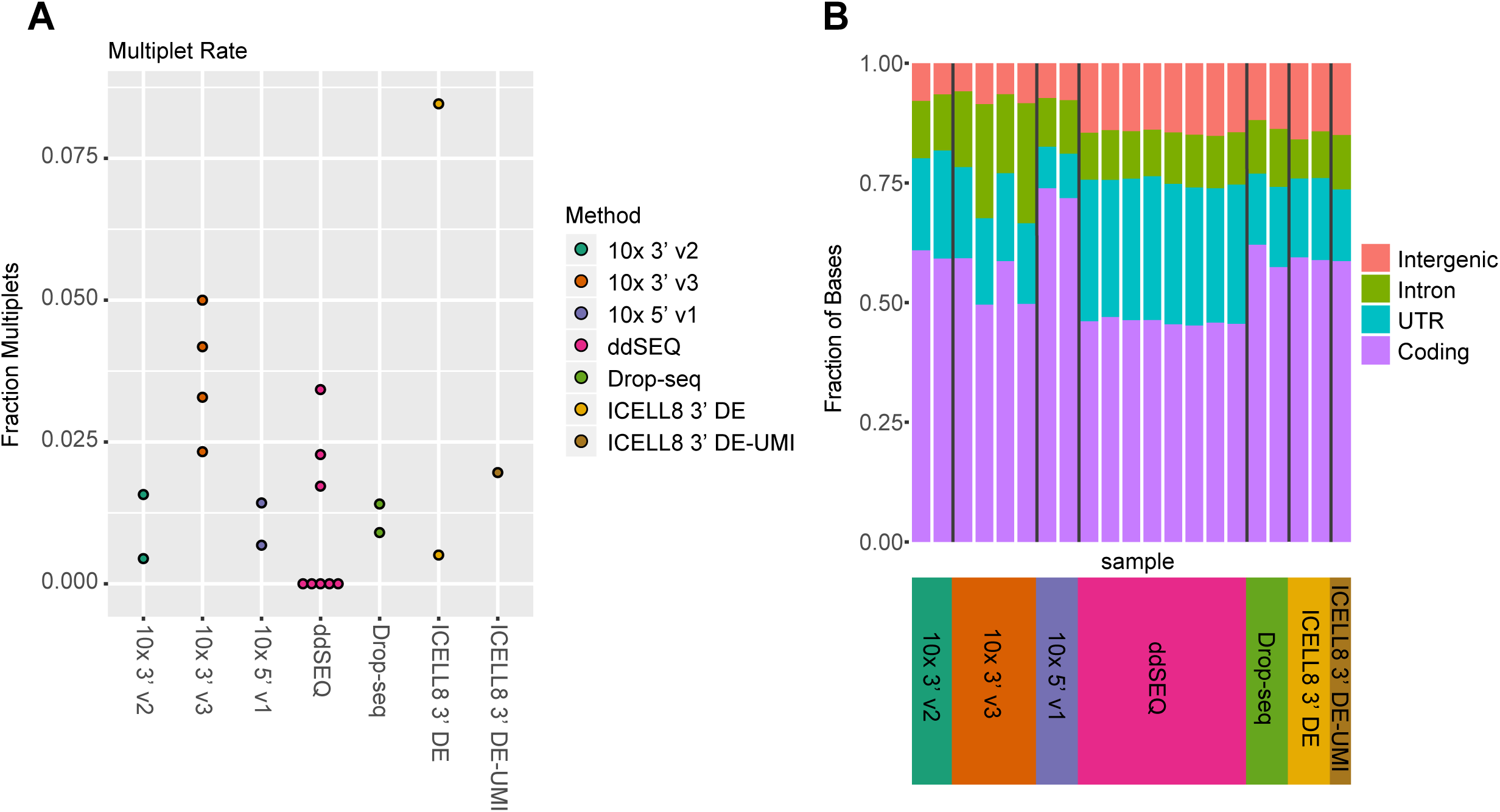
Library efficiencies. (A) Estimates of multiplet rates for each experiment based on number of CIDs with significant numbers of transcripts from human and mouse. Rates were adjusted to account for variable recovery of human and mouse cells. (B) Fraction of mapped bases aligning to intergenic (red), intronic (green), coding (purple), and untranslated (UTR) (cyan) regions by sample.

**Supplemental Figure 4:**
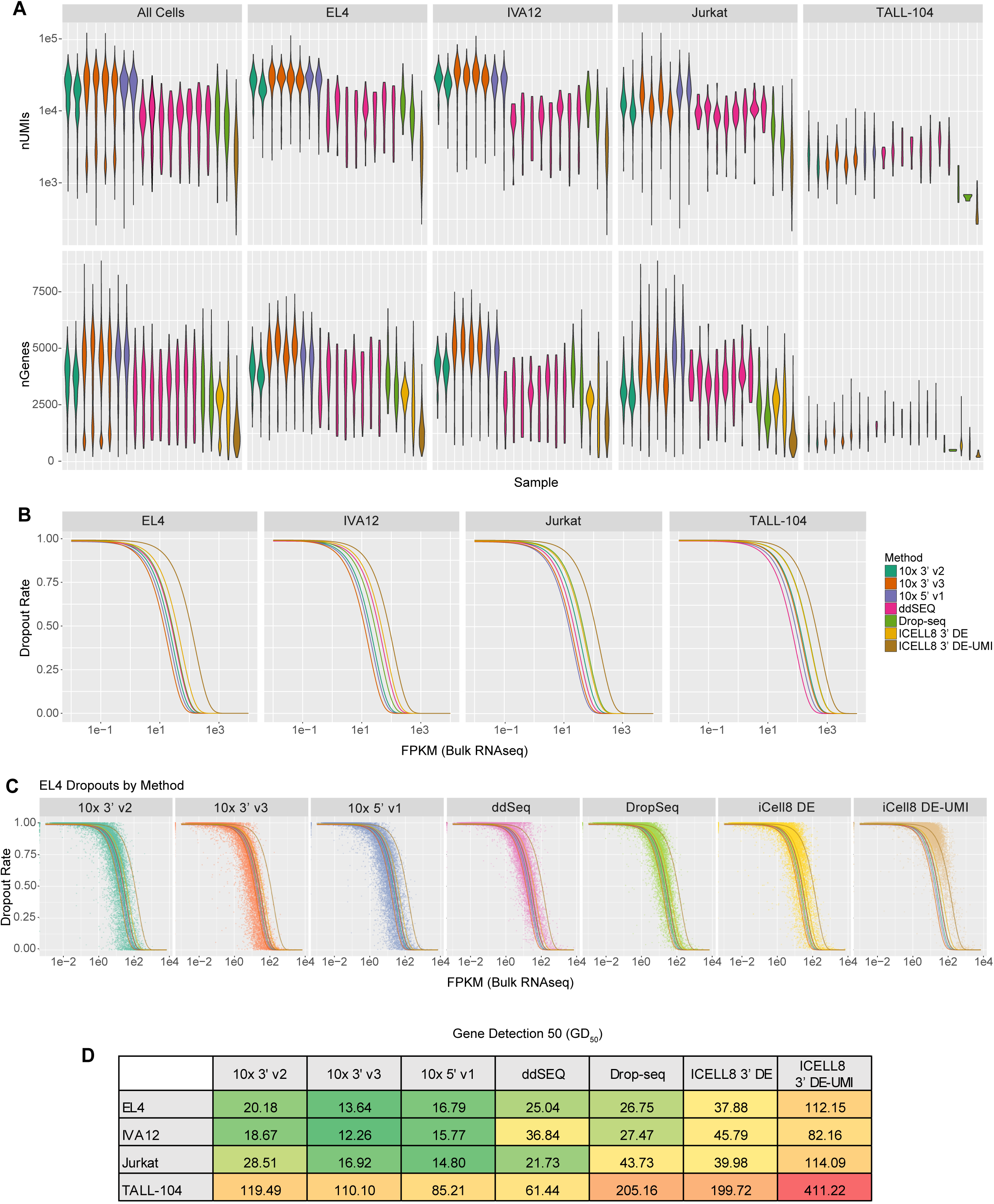
Transcript detection sensitivity. (A) Distributions of numbers of unique molecular identifiers (UMIs) and genes across all classified cells, or by cell-type. Read distribution was most consistent for EL4 cells across samples. (B) Models for dropout rate by expression level and cell type. A left shifted curve indicates higher sensitivity. (C) Dropout rates for mouse genes by expression level in bulk RNA-seq for EL4 cells. Solid lines indicate modeling curves for all methods. (D) GD_50_, the FPKM at which the dropout rate is expected to be 0.5, for dropout models by cell type. A low GD_50_ indicates high sensitivity.

**Supplemental Figure 5:**
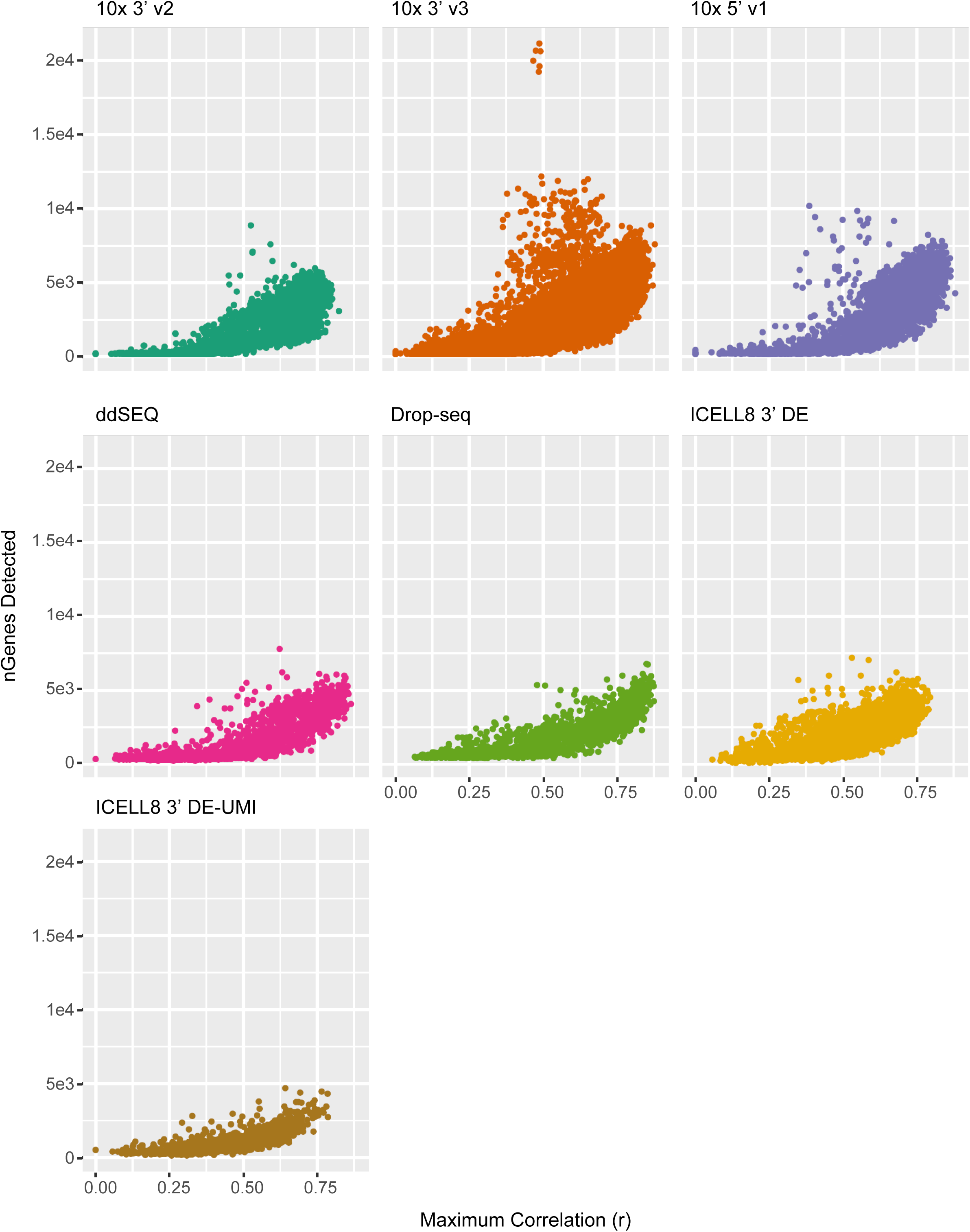
Correlation of single-cell RNA-seq to bulk RNA-seq. Number of detected genes were plotted as a function of the highest correlation coefficient *r* of CIDs above the inflection point to bulk RNA-seq.

**Supplemental Figure 6:**
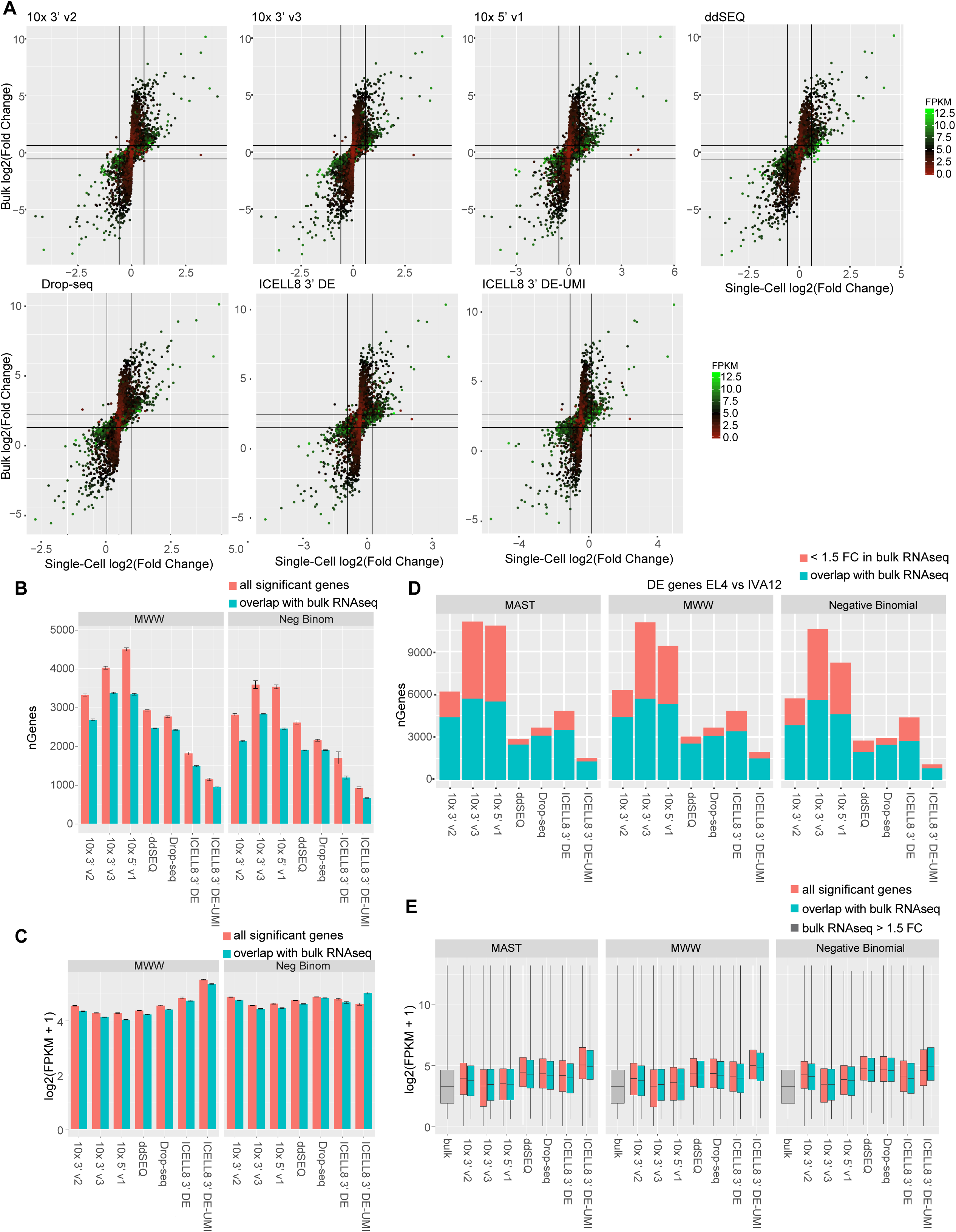
Differentially expressed (DE) genes. (A) Comparison of mouse gene fold changes in single-cell RNA-seq vs bulk RNA-seq for contrasting EL4 and IVA12 cells. Genes are colored by highest expression value (FPKM) in bulk RNA-seq data. Black lines indicate an absolute fold change of 1.5. While highly expressed genes in green correlate well between single-cell and bulk data, lowly expressed genes in red show little difference in expression in single-cell data. (B) Number of significant DE genes using the Mann-Whitney-Wilcoxon test or a negative binomial generalized linear model between EL4 and IVA12 cells. Error bars represent the 95% confidence interval from ten random sub-samplings of cells from each method. The total number of significant DE genes are plotted in red, the number of DE genes with > 1.5 fold difference in expression in bulk RNA-seq (5,868 genes) are plotted in cyan. (C) Median gene expression (FPKM) in bulk sequencing for all significant DE genes (red) or DE genes with > 1.5 fold difference in expression in bulk RNA-seq (cyan) are shown. Error bars represent 95% confidence interval. (D) Number of significant DE genes (FDR < 10^−4^) between all EL4 and IVA12 recovered cells for each method. Genes with > 1.5-fold change in bulk RNA-seq data are plotted in cyan with the remainder in red. (E) Distribution of gene expression level (FPKM) in bulk RNA-seq for DE genes identified using all cells for each method. All significant DE genes are plotted in red and DE genes with > 1.5-fold change in bulk RNA-seq data are plotted in cyan. Distribution of expression levels for all genes with > 1.5-fold change in bulk data are plotted in gray.

**Supplemental Table 1:** Summary of method designs.

| Method | Partitioning | Transcript Coverage | Bead Occupancy | Expected Cell Recovery | Cell Barcode and UMI [bp] relative to mRNA | Amplification Steps |
| --- | --- | --- | --- | --- | --- | --- |
| 10x Chromium Single Cell 3' v2 | Droplet | 3' end | 90% | 58% | 5'-Gene[10][16]-3' | Total cDNA + adapter PCR |
| 10x Chromium Single Cell 3' v3 | Droplet | 3' end | 90% | 62.50% | 5'-Gene[12][16]-3' | Total cDNA + adapter PCR |
| 10x Chromium Single Cell 5' v1 | Droplet | 5' end | 90% | 58% | 5'-[16][10]Gene-3' | Total cDNA + adapter PCR |
| Illumina/Bio-Rad ddSeq | Droplet | 3' end | 5% | 5% | 5'-Gene[8][3][6][15][6][15][6]-3' | Nextera PCR |
| Dolomite scRNA-Seq (Drop-seq) | Droplet | 3' end | Undisclosed | 3% | 5'-Gene[9][12]-3' | Total cDNA + Nextera PCR |
| Takara ICELL8 (3' DE, 3' DE-UMI) | Microwell | 3' end | N/A | 25% | 5'-Gene[14][11]-3' | Total cDNA + adapter PCR |

**Supplemental Table 2:** Sample summary.

| Sample ID | Method | Cell Mixture | EL4 % viable | IVA12 % viable | Jurkat % viable | TALL % viable | nCells Input |
| --- | --- | --- | --- | --- | --- | --- | --- |
| 1 | 10x 3' v2 | A | 93 | 95 | 96 | 50 | 8,700 |
| 2 | 10x 3' v2 | A | 93 | 95 | 96 | 50 | 3,500 |
| 21 | 10x 3' v3 | F | 98 | 93 | 99 | 65 | 4,800 |
| 22 | 10x 3' v3 | G | 98 | 93 | 99 | 65 | 4,800 |
| 23 | 10x 3' v3 | F | 98 | 93 | 99 | 65 | 9,600 |
| 24 | 10x 3' v3 | G | 98 | 93 | 99 | 65 | 9,600 |
| 5 | 10x 5' v1 | B | 89 | 92 | 99 | 89 | 8,700 |
| 6 | 10x 5' v1 | B | 89 | 92 | 99 | 89 | 3,500 |
| 9 | ddSEQ | C | 98 | 98 | 95 | 71 | 9,000 |
| 10 | ddSEQ | C | 98 | 98 | 95 | 71 | 9,000 |
| 11 | ddSEQ | C | 98 | 98 | 95 | 71 | 11,250 |
| 12 | ddSEQ | C | 98 | 98 | 95 | 71 | 11,250 |
| 13 | ddSEQ | D | 98 | 98 | 95 | 71 | 11,250 |
| 14 | ddSEQ | D | 98 | 98 | 95 | 71 | 11,250 |
| 15 | ddSEQ | D | 98 | 98 | 95 | 71 | 11,250 |
| 16 | ddSEQ | D | 98 | 98 | 95 | 71 | 11,250 |
| 19 | Drop-seq | E | 95 | 86 | 92 | 80 | 90,000 |
| 20 | Drop-seq | E | 95 | 86 | 92 | 80 | 180,000 |
| 27 | ICELL8 3' DE | H | 98 | 94 | 97 | 76 | 17,920 |
| 28 | ICELL8 3' DE | I | 95 | 63 | 97 | 70 | 17,920 |
| 29 | ICELL8 3' DE-UMI | I | 95 | 63 | 97 | 70 | 17,920 |

**Supplemental Table 3: Assessment of non-normalized scRNA-seq data (separate file)**. Sequenced reads were demultiplexed and aligned against a combined human and mouse reference genome with each technologies’ recommended software pipeline; all pipelines used a version of STAR for the alignment step. Aligned reads were MAPQ filtered and uniformly mapped against human and mouse gene annotations; read and UMI count matrices were derived using customized scripts. Cells and cell types were determined by similarity to bulk RNA-seq data.

**Supplemental Table 4:**
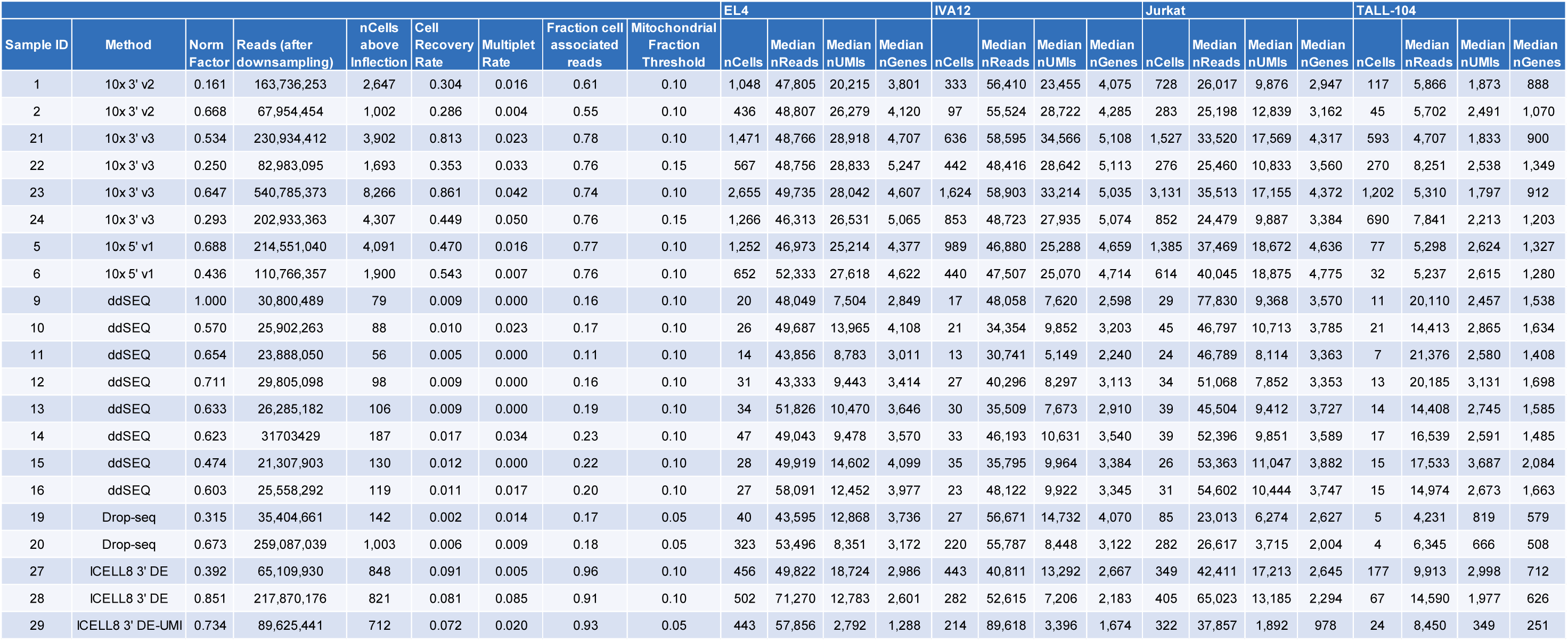
Results by sample.

**Supplemental Table 5:** Read-depth normalization iterations.

|  |  | EL4 |  |  |  |
| --- | --- | --- | --- | --- | --- |
| Sample ID | Method | nCells ± sd | median reads ± sd | median UMIs ± sd | median genes ± sd |
| 6 | 10x 5' v1 | 655.75 ± 2.63 | 52,280.88 ± 44.82 | 27,553.5 ± 43.04 | 4,613.25 ± 8.05 |
| 15 | ddSEQ | 27.5 ± 0.58 | 51,201 ± 1342.48 | 14,679.63 ± 64.47 | 4,106.88 ± 7.79 |
| 19 | Drop-seq | 40.5 ± 0.58 | 43,263.625 ± 254.09 | 12,676.63 ± 217.63 | 3,742 ± 9.97 |
| 27 | ICELL8 3' DE | 456.5 ± 0.58 | 49,828.63 ± 63.77 | 18,691.875 ± 60.74 | 2,981.75 ± 3.5 |
|  |  | IVA12 |  |  |  |
| Sample ID | Method | nCells ± sd | median reads ± sd | median UMIs ± sd | median genes ± sd |
| 6 | 10x 5' v1 | 438.5 ± 1.17 | 47,518.12 ± 19.8 | 25,095.63 ± 42.59 | 4,721.125 ± 9.52 |
| 15 | ddSEQ | 34.5 ± 1 | 36,343.5 ± 1113.78 | 10,017.25 ± 93.8 | 3,379.5 ± 16.34 |
| 19 | Drop-seq | 27 ± 0 | 56,397.75 ± 224.7 | 14,692 ± 84.7 | 4,062.25 ± 8.81 |
| 27 | ICELL8 3' DE | 443.75 ± 1.5 | 40,742.88 ± 120.14 | 13,242.75 ± 33.44 | 2,668.38 ± 2.06 |
|  |  | Jurkat |  |  |  |
| Sample ID | Method | nCells ± sd | median reads ± sd | median UMIs ± sd | median genes ± sd |
| 6 | 10x 5' v1 | 615.25 ± 1.5 | 40,019.88 ± 50.9 | 18,832.5 ± 33.45 | 4,773.63 ± 1.11 |
| 15 | ddSEQ | 26 ± 0 | 53,374.5 ± 109.5 | 11,027.25 ± 19.07 | 3,887.875 ± 11.78 |
| 19 | Drop-seq | 83 ± 1.83 | 23,522 ± 421.9 | 6,335.625 ± 56.45 | 2,634.38 ± 17.09 |
| 27 | ICELL8 3' DE | 348 ± 1.41 | 42,004.25 ± 363.66 | 17,143.25 ± 69.09 | 2,648.625 ± 7.54 |
|  |  | Tall |  |  |  |
| Sample ID | Method | nCells ± sd | median reads ± sd | median UMIs ± sd | median genes ± sd |
| 6 | 10x 5' v1 | 32.75 ± 1.5 | 5,224.5 ± 83.5 | 2,485.63 ± 90.57 | 1,257.88 ± 20.95 |
| 15 | ddSEQ | 14.75 ± 0.5 | 17,661.875 ± 267.97 | 3,747.25 ± 127.94 | 2,086.75 ± 27.07 |
| 19 | Drop-seq | 5 ± 0 | 4,201.25 ± 30.23 | 816.75 ± 7.41 | 581.75 ± 4.86 |
| 27 | ICELL8 3' DE | 177 ± 0.82 | 10,029.75 ± 86.96 | 2,986.125 ± 23.48 | 711.25 ± 0.5 |

## References

1. Bandura DR, Baranov VI, Ornatsky OI, Antonov A, Kinach R, Lou X, et al. Mass cytometry: technique for real time single cell multitarget immunoassay based on inductively coupled plasma time-of-flight mass spectrometry. Anal Chem. 2009;81(16):6813–22. doi: 10.1021/ac901049w. PubMed PMID: 19601617.

2. He H, Suryawanshi H, Morozov P, Gay-Mimbrera J, Del Duca E, Kim HJ, et al. Single-cell transcriptome analysis of human skin identifies novel fibroblast subpopulation and enrichment of immune subsets in atopic dermatitis. J Allergy Clin Immunol. 2020. doi: 10.1016/j.jaci.2020.01.042. PubMed PMID: 32035984.

3. Sade-Feldman M, Yizhak K, Bjorgaard SL, Ray JP, de Boer CG, Jenkins RW, et al. Defining T Cell States Associated with Response to Checkpoint Immunotherapy in Melanoma. Cell. 2018;175(4):998–1013 e20. doi: 10.1016/j.cell.2018.10.038. PubMed PMID: 30388456; PubMed Central PMCID: PMCPMC6641984.

4. Zhang Y, Zheng L, Zhang L, Hu X, Ren X, Zhang Z. Deep single-cell RNA sequencing data of individual T cells from treatment-naive colorectal cancer patients. Sci Data. 2019;6(1):131. doi: 10.1038/s41597-019-0131-5. PubMed PMID: 31341169; PubMed Central PMCID: PMCPMC6656756.

5. Zheng C, Zheng L, Yoo JK, Guo H, Zhang Y, Guo X, et al. Landscape of Infiltrating T Cells in Liver Cancer Revealed by Single-Cell Sequencing. Cell. 2017;169(7):1342–56 e16. doi: 10.1016/j.cell.2017.05.035. PubMed PMID: 28622514.

6. Zilionis R, Engblom C, Pfirschke C, Savova V, Zemmour D, Saatcioglu HD, et al. Single-Cell Transcriptomics of Human and Mouse Lung Cancers Reveals Conserved Myeloid Populations across Individuals and Species. Immunity. 2019;50(5):1317–34 e10. doi: 10.1016/j.immuni.2019.03.009. PubMed PMID: 30979687; PubMed Central PMCID: PMCPMC6620049.

7. Zhang L, Li Z, Skrzypczynska KM, Fang Q, Zhang W, O’Brien SA, et al. Single-Cell Analyses Inform Mechanisms of Myeloid-Targeted Therapies in Colon Cancer. Cell. 2020;181(2):442-59 e29. Epub 2020/04/18. doi: 10.1016/j.cell.2020.03.048. PubMed PMID: 32302573.

8. Ziegenhain C, Vieth B, Parekh S, Reinius B, Guillaumet-Adkins A, Smets M, et al. Comparative Analysis of Single-Cell RNA Sequencing Methods. Mol Cell. 2017;65(4):631–43 e4. doi: 10.1016/j.molcel.2017.01.023. PubMed PMID: 28212749.

9. Svensson V, Natarajan KN, Ly LH, Miragaia RJ, Labalette C, Macaulay IC, et al. Power analysis of single-cell RNA-sequencing experiments. Nat Methods. 2017;14(4):381–7. doi: 10.1038/nmeth.4220. PubMed PMID: 28263961; PubMed Central PMCID: PMCPMC5376499.

10. Mereu E, Lafzi A, Moutinho C, Ziegenhain C, McCarthy DJ, Álvarez-Varela A, et al. Benchmarking single-cell RNA-sequencing protocols for cell atlas projects. Nature Biotechnology. 2020. doi: 10.1038/s41587-020-0469-4.

11. Tian L, Dong X, Freytag S, Le Cao KA, Su S, JalalAbadi A, et al. Benchmarking single cell RNA-sequencing analysis pipelines using mixture control experiments. Nat Methods. 2019;16(6):479–87. doi: 10.1038/s41592-019-0425-8. PubMed PMID: 31133762.

12. Zhang X, Li T, Liu F, Chen Y, Yao J, Li Z, et al. Comparative Analysis of Droplet-Based Ultra-High-Throughput Single-Cell RNA-Seq Systems. Mol Cell. 2019;73(1):130–42 e5. doi: 10.1016/j.molcel.2018.10.020. PubMed PMID: 30472192.

13. Ding J, Adiconis X, Simmons SK, Kowalczyk MS, Hession CC, Marjanovic ND, et al. Systematic comparison of single-cell and single-nucleus RNA-sequencing methods. Nat Biotechnol. 2020. Epub 2020/04/29. doi: 10.1038/s41587-020-0465-8. PubMed PMID: 32341560.

14. Zheng GX, Terry JM, Belgrader P, Ryvkin P, Bent ZW, Wilson R, et al. Massively parallel digital transcriptional profiling of single cells. Nat Commun. 2017;8:14049. doi: 10.1038/ncomms14049. PubMed PMID: 28091601; PubMed Central PMCID: PMCPMC5241818

15. L.M., D.A.M., S.Y.N., M.S.L., P.W.W., C.M.H., R.B., A.W., K.D.N., T.S.M. and B.J.H. are employees of 10x Genomics. Macosko EZ, Basu A, Satija R, Nemesh J, Shekhar K, Goldman M, et al. Highly Parallel Genome-wide Expression Profiling of Individual Cells Using Nanoliter Droplets. Cell. 2015;161(5):1202–14. doi: 10.1016/j.cell.2015.05.002. PubMed PMID: 26000488; PubMed Central PMCID: PMCPMC4481139.

16. Goldstein LD, Chen YJ, Dunne J, Mir A, Hubschle H, Guillory J, et al. Massively parallel nanowell-based single-cell gene expression profiling. BMC Genomics. 2017;18(1):519. doi: 10.1186/s12864-017-3893-1. PubMed PMID: 28687070; PubMed Central PMCID: PMCPMC5501953.

17. Picelli S, Bjorklund AK, Faridani OR, Sagasser S, Winberg G, Sandberg R. Smart-seq2 for sensitive full-length transcriptome profiling in single cells. Nat Methods. 2013;10(11):1096–8. doi: 10.1038/nmeth.2639. PubMed PMID: 24056875.

18. Picelli S, Faridani OR, Bjorklund AK, Winberg G, Sagasser S, Sandberg R. Full-length RNA-seq from single cells using Smart-seq2. Nat Protoc. 2014;9(1):171–81. doi: 10.1038/nprot.2014.006. PubMed PMID: 24385147.

19. Islam S, Zeisel A, Joost S, La Manno G, Zajac P, Kasper M, et al. Quantitative single-cell RNA-seq with unique molecular identifiers. Nat Methods. 2014;11(2):163–6. doi: 10.1038/nmeth.2772. PubMed PMID: 24363023.

20. Andrews TS, Hemberg M. M3Drop: dropout-based feature selection for scRNASeq. Bioinformatics. 2019;35(16):2865–7. doi: 10.1093/bioinformatics/bty1044. PubMed PMID: 30590489; PubMed Central PMCID: PMCPMC6691329.

21. Tunnacliffe E, Chubb JR. What Is a Transcriptional Burst? Trends Genet. 2020. doi: 10.1016/j.tig.2020.01.003. PubMed PMID: 32035656.

22. Lenstra TL, Rodriguez J, Chen H, Larson DR. Transcription Dynamics in Living Cells. Annu Rev Biophys. 2016;45:25–47. doi: 10.1146/annurev-biophys-062215-010838. PubMed PMID: 27145880; PubMed Central PMCID: PMCPMC6300980.

23. Finak G, McDavid A, Yajima M, Deng J, Gersuk V, Shalek AK, et al. MAST: a flexible statistical framework for assessing transcriptional changes and characterizing heterogeneity in single-cell RNA sequencing data. Genome Biol. 2015;16:278. doi: 10.1186/s13059-015-0844-5. PubMed PMID: 26653891; PubMed Central PMCID: PMCPMC4676162.

24. Soneson C, Robinson MD. Bias, robustness and scalability in single-cell differential expression analysis. Nat Methods. 2018;15(4):255-61. Epub 2018/02/27. doi: 10.1038/nmeth.4612. PubMed PMID: 29481549.

25. Stuart T, Butler A, Hoffman P, Hafemeister C, Papalexi E, Mauck WM, 3rd, et al. Comprehensive Integration of Single-Cell Data. Cell. 2019;177(7):1888-902 e21. Epub 2019/06/11. doi: 10.1016/j.cell.2019.05.031. PubMed PMID: 31178118; PubMed Central PMCID: PMCPMC6687398.

26. Lun ATL, Riesenfeld S, Andrews T, Dao TP, Gomes T, participants in the 1st Human Cell Atlas J, et al. EmptyDrops: distinguishing cells from empty droplets in droplet-based single-cell RNA sequencing data. Genome Biol. 2019;20(1):63. doi: 10.1186/s13059-019-1662-y. PubMed PMID: 30902100; PubMed Central PMCID: PMCPMC6431044.

27. Wang YJ, Schug J, Lin J, Wang Z, Kossenkov A,, et al. Comparative analysis of commercially available single-cell RNA sequencing platforms for their performance in complex human tissues. bioRxiv. 2019. doi: https://doi.org/10.1101/541433.

28. Lu DR, Wu H, Driver I, Ingersoll S, Sohn S, Wang S, et al. Dynamic changes in the regulatory T-cell heterogeneity and function by murine IL-2 mutein. Life Sci Alliance. 2020;3(5). Epub 2020/04/10. doi: 10.26508/lsa.201900520. PubMed PMID: 32269069; PubMed Central PMCID: PMCPMC7156283.

29. Hanson WM, Chen Z, Jackson LK, Attaf M, Sewell AK, Heemstra JM, et al. Reversible Oligonucleotide Chain Blocking Enables Bead Capture and Amplification of T-Cell Receptor alpha and beta Chain mRNAs. J Am Chem Soc. 2016;138(35):11073-6. Epub 2016/08/02. doi: 10.1021/jacs.6b04465. PubMed PMID: 27478996; PubMed Central PMCID: PMCPMC5249220.

30. Saikia M, Burnham P, Keshavjee SH, Wang MFZ, Heyang M, Moral-Lopez P, et al. Simultaneous multiplexed amplicon sequencing and transcriptome profiling in single cells. Nat Methods. 2019;16(1):59-62. Epub 2018/12/19. doi: 10.1038/s41592-018-0259-9. PubMed PMID: 30559431; PubMed Central PMCID: PMCPMC6378878.

31. Liao Y, Smyth GK, Shi W. featureCounts: an efficient general purpose program for assigning sequence reads to genomic features. Bioinformatics. 2014;30(7):923-30. Epub 2013/11/15. doi: 10.1093/bioinformatics/btt656. PubMed PMID: 24227677.

32. Dobin A, Davis CA, Schlesinger F, Drenkow J, Zaleski C, Jha S, et al. STAR: ultrafast universal RNA-seq aligner. Bioinformatics. 2013;29(1):15-21. Epub 2012/10/30. doi: 10.1093/bioinformatics/bts635. PubMed PMID: 23104886; PubMed Central PMCID: PMCPMC3530905.

33. Romagnoli D, Boccalini G, Bonechi M, Biagioni C, Fassan P, Bertorelli R, et al. ddSeeker: a tool for processing Bio-Rad ddSEQ single cell RNA-seq data. BMC Genomics. 2018;19(1):960. Epub 2018/12/26. doi: 10.1186/s12864-018-5249-x. PubMed PMID: 30583719; PubMed Central PMCID: PMCPMC6304778.

34. Li H, Handsaker B, Wysoker A, Fennell T, Ruan J, Homer N, et al. The Sequence Alignment/Map format and SAMtools. Bioinformatics. 2009;25(16):2078-9. Epub 2009/06/10. doi: 10.1093/bioinformatics/btp352. PubMed PMID: 19505943; PubMed Central PMCID: PMCPMC2723002.

